# Association of bidirectional network cores in the brain with perceptual awareness and cognition

**DOI:** 10.1101/2024.04.30.591001

**Authors:** Tomoya Taguchi, Jun Kitazono, Shuntaro Sasai, Masafumi Oizumi

## Abstract

The brain comprises a complex network of interacting regions. To understand the roles and mechanisms of this intricate network, it is crucial to elucidate its structural features related to cognitive functions. Recent empirical evidence suggests that both feedforward and feedback signals are necessary for conscious perception, emphasizing the importance of subnetworks with bidirectional interactions. However, the link between such subnetworks and conscious perception remains unclear due to the complexity of brain networks. In this study, we propose a framework for extracting subnetworks with strong bidirectional interactions—termed the “cores” of a network—from brain activity. We applied this framework to resting-state and task-based human fMRI data from participants of both sexes to identify regions forming strongly bidirectional cores. We then explored the association of these cores with conscious perception and cognitive functions. We found that the extracted central cores predominantly included cerebral cortical regions rather than subcortical regions. Additionally, regarding their relation to conscious perception, we demonstrated that the cores tend to include regions previously reported to be affected by electrical stimulation that altered conscious perception, although the results are not statistically robust due to the small sample size. Furthermore, in relation to cognitive functions, based on a meta-analysis and comparison of the core structure with a cortical functional connectivity gradient, we found that the central cores were related to unimodal sensorimotor functions. The proposed framework provides novel insights into the roles of network cores with strong bidirectional interactions in conscious perception and unimodal sensorimotor functions.

**Significance Statement:** To understand the brain’s network, we need to decipher its structural features linked to cognitive functions. Recent studies suggest the importance of subnetworks with bidirectional interactions for conscious perception, but their exact relationship remains unclear due to the brain’s complexity. Here we propose a framework for extracting subnetworks with strong bidirectional interactions, or network “cores.” We applied it to fMRI data and explored the association of the cores with conscious perception and cognitive functions. The central cores predominantly included cortical regions rather than subcortical ones, and tended to comprise previously reported regions wherein electrical stimulation altered perception, suggesting the potential importance of bidirectional cores for conscious perception. Additionally, further analysis revealed the relationship of the cores to unimodal sensorimotor functions.

## Introduction

Complex networks formed by interactions between brain regions are involved in cognitive functions. Analyzing these networks using network theory could crucially enable the identification of network features relevant to specific cognitive functions, which may further our understanding of the mechanisms that underlie cognitive functions (Liao et al., 2017; Bullmore and Sporns, 2009; Sporns, 2018). For example, studies have shown that the manifestation of module structures (i.e., modularity) predicts high performance in working memory tasks (Stevens et al., 2012; Finc et al., 2017; Gamboa et al., 2014; Liang et al., 2016; Finc et al., 2020). Moreover, research on small-world properties has demonstrated that the efficiency of signal propagation between brain regions (i.e., network efficiency) can predict an individual’s intellectual capacity (van den Heuvel et al., 2009).

One relationship between network features and cognitive functions that has attracted considerable attention is the link between the bidirectionality of networks and conscious perception. Empirical research has shown that conscious perception of a sensory stimulus requires bidirectional signaling that involves both feedforward and feedback signal propagation (Cauller and Kulics, 1991; Lamme et al., 1998; Supèr et al., 2001; Self et al., 2012; Auksztulewicz et al., 2012; Sachidhanandam et al., 2013; Tang et al., 2014; Koivisto et al., 2014; Manita et al., 2015). Moreover, the importance of bidirectional interactions in consciousness is independent of sensory modality (Dembski et al., 2021) (vision (Lamme et al., 1998; Supèr et al., 2001; Self et al., 2012; Tang et al., 2014; Koivisto et al., 2014), somatosensation (Cauller and Kulics, 1991; Auksztulewicz et al., 2012; Sachidhanandam et al., 2013; Manita et al., 2015), and audition (Gutschalk et al., 2008; Dykstra et al., 2016; Eklund and Wiens, 2019; Schlossmacher et al., 2021; Hayat et al., 2022)).

Based on these experimental findings and theoretical insights, to understand the relationship between the brain network and conscious perception, the identification of brain subnetworks that have strong bidirectional interactions and ascertaining how these subnetworks relate to conscious perception is pivotal. However, the identification of such subnetworks has been challenging. To date, methods for extracting subnetworks with strong connections have been applied to brain functional networks (e.g., s-core decomposition (Chatterjee and Sinha, 2007; van den Heuvel and Sporns, 2011; Harriger et al., 2012; Crobe et al., 2016), network hubs (van den Heuvel and Sporns, 2013; Royer et al., 2022), rich clubs (van den Heuvel and Sporns, 2011; Liang et al., 2018; Wang et al., 2020), and modularity (Bertolero et al., 2015; Chen et al., 2021)). However, the abovementioned methods do not consider the direction of influence, particularly the bidirectionality of interactions. Consequently, the locations of the subnetworks with pronounced bidirectionality and their links to cognitive functions, including conscious perception, remain elusive.

To address this gap, we propose a novel framework for extracting subnetworks with strong bidirectional interactions from brain activity that were designated as “cores.” We applied this framework to functional magnetic resonance imaging (fMRI) data to identify the regions that constitute strongly bidirectional cores and subsequently performed two analyses on the identified regions: one that examined the relationship of the core regions with conscious perception and the other that explored the association of the core regions with cognitive functions. The following section provides an overview of this analysis.

First, we investigated the brain regions that were more likely to be components of strongly bidirectional cores. If certain brain regions consistently appear in the cores under various conditions, this would suggest their general importance in conscious perception and a broad range of cognitive functions. Therefore, using data from the Human Connectome Project (HCP) (Van Essen et al., 2013), we extracted the cores at rest and during seven cognitive tasks. Our objective was to determine the brain regions that consistently form strongly bidirectional cores under resting-state and task conditions.

We subsequently investigated the association of the core regions with conscious perception. We focused on a previous study wherein electrical brain stimulation was used to assess the association of brain regions with conscious perception (Fox et al., 2020). We posited that, if regions within strongly bidirectional cores are integral to conscious perception, then, stimulation of these regions would propagate effects throughout the cores, and thereby result in alterations of conscious perception. Therefore, in this study, we compared the likelihood of each region of interest (ROI) being included in strongly bidirectional cores and the rate at which the intracranial electrical stimulation (iES) (Fox et al., 2020) of each ROI elicited a change in conscious perception. iES involves direct electrical stimulation of the brain via intracranially placed electrodes (Borchers et al., 2011) that enable causal modulation of neural activity. Prior research has quantitatively evaluated the extent to which the iES of each brain region induces reportable changes in conscious perceptual experiences across the entire cerebral cortex (Fox et al., 2020). We investigated whether the rates of perceptual change elicited by iES (elicitation rates) were associated with the probability of being included in strongly bidirectional cores. Higher elicitation rates of regions that are more likely to be part of strong cores would suggest a significant association between strongly bidirectional cores and conscious perception.

Then, to investigate the association of the cores with cognitive functions more broadly, we performed a hypothesis-free meta-analysis using NeuroSynth (Yarkoni et al., 2011). NeuroSynth is a platform for examining associations between brain regions and specific psychological and neurological terms based on a large database of fMRI studies. Using this tool, thousands of fMRI data can be aggregated to identify statistical associations between ROIs and specific terms, such as “memory” and “attention.” This approach facilitated the identification of differences in the associated terms between ROIs with a strong or weak tendency to be included in the cores. Finally, to further assess the association of the cores with cognitive function, we examined the relationship between the cores to a functional connectivity gradient (Margulies et al., 2016). The gradient is known to correspond to a spatial gradient over the cerebral cortex, ranging from unimodal sensory to higher-order association regions.

## Materials and Methods

### fMRI Data Acquisition and Preprocessing

The 3T-fMRI data were obtained from the Washington University-Minnesota Consortium Human Connectome Project (HCP, (Van Essen et al., 2013)). The data were approved by the Institutional Review Board of the University of Washington. From the HCP 1200 Subjects Data Release (Van Essen et al., 2013) (www.humanconnectome.org/), we used 352 subjects in accordance with the criteria proposed in ref. (Ito et al., 2020). The selection of the 352 participants (183 females, ages 22-36 years) was based on three main criteria: quality control assessments, motion artifacts, and family relations. Participants were excluded if they had any quality control flags, including focal anatomical anomalies found in T1-weighted or T2-weighted scans, focal segmentation or surface errors detected by the HCP structural pipeline, data collected during periods with known issues with the head coil, or data where some of the FIX-ICA components were manually reclassified. Participants were also excluded if any fMRI run had more than 50% of the repetition times (TRs) with framewise displacement greater than 0.25 mm to minimize motion artifacts. Additionally, only unrelated participants (i.e., non-familial) were included, and those without genotype testing data were excluded. A full list of participants are included as part of the code release of ref. (Ito et al., 2020).

In the HCP study, imaging was performed on a 3T Siemens Skyra scanner with a 32-channel head coil, utilizing a multiband acceleration factor of 8. The imaging parameters included a TR of 720 ms, TE of 33.1 ms, a flip angle of 52°, and an isotropic resolution of 2.0 mm. The data collection spanned two days: Day 1 involved anatomical scans (T1 and T2-weighted images at 0.7 mm isotropic resolution), followed by two 14.4-minute resting-state fMRI scans (Rest 1, the left-right (LR) and right-left (RL) scans) and task fMRI scans. The second day included a diffusion imaging scan, another two 14.4-minute resting-state fMRI scans (Rest 2, LR and RL scans), and additional task fMRI sessions.

We started with minimally preprocessed fMRI data at rest and during seven cognitive tasks (emotion, gambling, language, motor, relational, social, and working memory), and we performed denoising by estimating nuisance regressors and subtracting them from the signal at every vertex (Satterthwaite et al., 2013). Accordingly, we used 36 nuisance and spike regressors, developed by ref. (Satterthwaite et al., 2013), that comprised: (1–6) six motion parameters, (7) white-matter time-series, (8) cerebrospinal fluid time-series, (9) global signal time-series, (10–18) temporal derivatives of (1–9), and (19–36) quadratic terms for (1–18). The spike regressors were computed as described previously in ref. (Satterthwaite et al., 2013), with 1.5-mm movement used as a spike-identification threshold. These fMRI data have a temporal resolution (TR) of 0.72 seconds. The number of frames for rest and each task state is as follows: emotion, 176; gambling, 253; language, 316; motor, 284; relational, 232; social, 274; working memory, 405; and rest, 1200 frames.

Using the methodology proposed in ref. (Luppi and Stamatakis, 2021), we parcellated the cerebral cortex and subcortex into 200 and 32 ROIs, based on Schaefer’s parcellation (Schaefer et al., 2018) and Tian’s parcellation (Tian et al., 2020), respectively, that generated a total of 232 ROIs. Functional and structural networks obtained using this parcellation are close to the average of those derived from other parcellations, which makes them representative networks (Luppi and Stamatakis, 2021) and, accordingly, we employed this parcellation in this study. We normalized the time-series data for each session to z-scores and then applied a 0.008–0.08 Hz secondorder Butterworth band-pass filter.

### Estimation of directed functional network

To extract cores with strong bidirectional interactions, it is first necessary to quantify the statistical causal strengths between brain regions and construct a directed network. For this purpose, we computed the normalized directed transfer entropy (NDTE, (Deco et al., 2021)) for each pair of ROIs from the BOLD signals: We fitted a bivariate vector autoregressive (VAR) model to the BOLD signals *X* and *Y* of each pair of ROIs and set the lag order *T* of the VAR model to 10. A bivariate VAR(*T*) model for the process 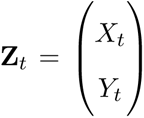, where **Z***_t_* is a 2-dimensional vector of the two time series at time *t*, takes the form:

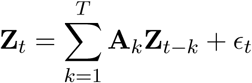

where **A***_k_* are 2×2 regression coefficient matrices, and the 2-dimensional vector *ɛ_t_* is the residuals. The parameters of the model are the coefficient matrices **A***_k_* and the covariance matrix of the residuals Σ = cov(*ɛ_t_*), which is time-invariant due to stationarity.

The estimation of the VAR model was performed using locally weighted regression, using data from the left-right (LR) and right-left (RL) scans of 352 subjects as trials, resulting in 704 trials for each graph of each task (emotion, gambling, language, motor, relational, social, and working memory) and rest (Rest 1, Rest 2). There were two resting-state sessions in the HCP data, Rest 1 and Rest 2, and here we constructed graphs for each of the Rest 1 and Rest 2 sessions. The Multivariate Granger Causality Toolbox (Barnett and Seth, 2014) was employed for these computations.

Subsequently, the transfer entropy (TE) for each pair of ROIs was calculated using the parameters of this estimated VAR model. TE *T*_XY_ from the ROI X to ROI Y quantifies how much the past activity of X contributes to predicting the future of Y (Deco et al., 2021). Mathematically, *T*_XY_ is conditional mutual information *I* (*Y_i_*_+1_; *X^i^* | *Y^i^*) that can be written using entropy as

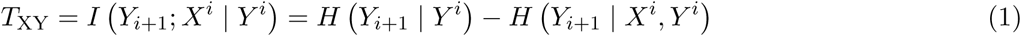

where *Y_i_*_+1_ denotes the activity of the ROI Y at time *i* + 1, and *X^i^* and *Y^i^* denote the history of activity of X and Y (*X^i^* := (*X_i_, X_i_*_−1_*,…, X_i_*_−(_*_T_* _−1)_) and *Y^i^* := (*Y_i_, Y_i_*_−1_*,…, Y_i_*_−(_*_T_* _−1)_), respectively). Thus, TE quantifies the reduction in uncertainty of *Y_i_*_+1_ when *Y^i^* and *X^i^* are known, compared to when only *Y^i^* is known. Then, aiming to add and compare the TE between different ROI pairs, we normalized TE by mutual information and obtained NDTE *F*_XY_ (Deco et al., 2021):

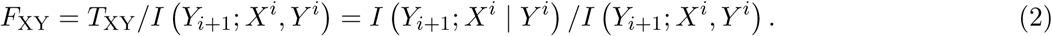

The normalization factor *I* (*Y_i_*_+1_; *X^i^, Y^i^*) is the mutual information between *Y_i_*_+1_ and *X^i^, Y^i^*, and can be decomposed as

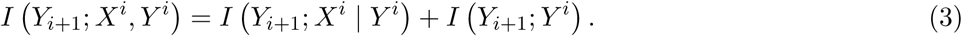

This decomposition indicates that the normalization compares the predictability of *Y_i_*_+1_ by *X^i^* | *Y^i^* (i.e., *I* (*Y_i_*_+1_; *X^i^* | *Y^i^*)) with the internal predictability of *Y_i_*_+1_ by *Y^i^* (i.e., *I* (*Y_i_*_+1_; *Y^i^*)).

First, we performed edge thresholding based on statistical significance. To achieve this, we generated 100 surrogate data sets using block permutation (via block permute 01 from the MVGC toolbox) and applied kernel density estimation (from the MATLAB function ksdensity) to their distributions to calculate the p-values. For the block size in block permutation, we followed the recommendation of setting it to the order of the VAR model (Barnett and Seth, 2014) and used a block size of 10. After obtaining the p-values, we retained only the edges that surpassed the significance level (*α*= 0.05, with Bonferroni correction).

As mentioned above, we used the block permutation method to generate surrogate data, whereas the original NDTE framework used the circular shift method (Deco et al., 2021). The reason we did not use the circular shift method is that the method is not suitable for data with a short length. The circular shift method generates only a limited number of surrogates when data length is short. For example, for the Emotion task, which contains approximately 200 time points, if we set the minimum shift width to 10 time points, as done in the study by Deco et al., we obtain only about 20 surrogates that differ from each other by more than 10 time points. This increases the likelihood that the surrogates will closely resemble each other if the number of surrogates is more than 20. In contrast for the block permutation, with a block size of 10, as we did, it divides the 200 time points into 20 blocks of 10 points each, yielding approximately 20! unique combinations. This approach substantially increases the number of possible surrogates and reduces the chance of surrogates resembling each other, making block permutation a more robust choice for surrogate data generation.

Subsequently, we adjusted the graphs to have a fixed density, as it is generally considered appropriate to compare graphs with equal densities (van Wijk et al., 2010; Luppi and Stamatakis, 2021; van den Heuvel et al., 2017; Jalili, 2016). Given that anatomical connections in brain networks are typically sparse (with densities around 20% or less) (Bullmore and Sporns, 2012; Hagmann et al., 2008; Luppi and Stamatakis, 2021), and that much of the functional connectivity may be spurious due to this sparsity (Luppi and Stamatakis, 2021), we retained only the strongest edges to match the reference densities of 5%, 10%, and 20%.

### Extraction of bidirectionally interacting cores

From the directed network constructed using NDTE, we extracted subnetworks with particularly strong bidirectional connections—the network “cores.” To identify such cores, our proposed framework uses the algorithm proposed by Kitazono et al. (Kitazono et al., 2023). This algorithm enables the hierarchical decomposition of the entire network based on the strength of bidirectional connections.

To demonstrate how the algorithm works, we present its application to a simple toy network in Fig. **1c**. In the toy network, the node set {*B, E, F, I, J*} is connected bidirectionally whereas the node set {*A, C, D, G, H*} is connected in a feedforward manner. Extracting cores from this network according to the method of Kitazono et al. (Kitazono et al., 2023) reveals the network cores, as indicated by various shades of color. The subnetwork {*E, F, I, J*}, highlighted in orange, represents the core with the strongest bidirectional connections, followed by the subnetwork {*B, E, F, I, J*}, shown in blue. As shown in this example, cores with stronger bidirectional connections are nested within those with weaker bidirectional connections and thus generally form a unimodal or multimodal hierarchical structure.

**Figure 1:**
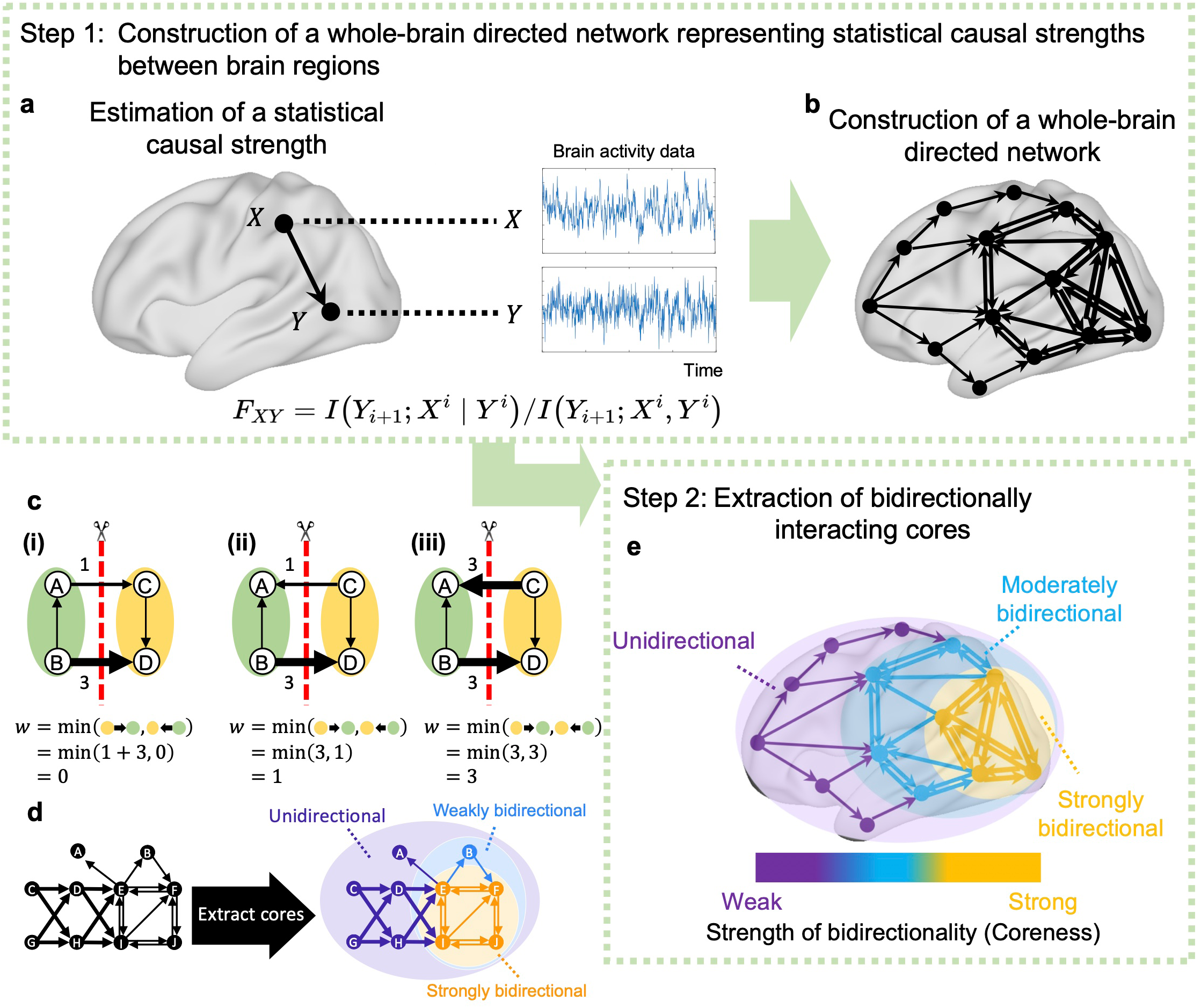
Schematic of our framework for extracting bidirectionally interacting cores from a whole-brain directed network. Our proposed framework in this study comprises two steps. In the first step, based on brain activity, we estimate the strength of the statistical causal influence between brain regions. a, Specifically, we calculate the NDTE *F*_XY_ (Deco et al., 2021) from the brain activities of regions X and Y. NDTE *F*_XY_ represents the strength of statistical causal influence from the source brain region X to the target brain region Y (See Methods for details). **b**, We then apply this approach to all pairs of brain regions across the whole brain and thereby construct a whole-brain-directed network. **c**, Examples of the strength of bidirectional connections. **i**, When two parts of a network are linked unidirectionally, the strength of bidirectional connections *w*(*V*_L_; *V*_R_) equals 0. **ii**, When a connection in one direction is strong (*w*(*V*_L_ → *V*_R_) = 3), while the connection in the other direction is weak (*w*(*V*_R_ → *V*_L_) = 1), the strength of bidirectional connections is small (*w*(*V*_L_; *V*_R_) = 1). **iii**, When connections in both directions are strong (*w*(*V*_L_ → *V*_R_) = *w*(*V*_R_ → *V*_L_) = 3), the strength of bidirectional connections is large (*w*(*V*_L_; *V*_R_) = 3). **d**, An example of complexes using a toy network. Nodes BEFIJ are connected bidirectionally whereas nodes ACDGH are connected in a feedforward manner. Strongly bidirectional cores in this network are indicated by a colored background, wherein subnet EFIJ, in orange, has the strongest bidirectional connections, followed by BEFIJ, in blue. Generally, complexes with stronger bidirectional connections are included in those with weaker bidirectional connections to form a unimodal or multimodal hierarchical structure. **e**, In the second step, from the constructed directed network, we extract cores with strong bidirectional connections. To extract cores, we use the algorithm by Kitazono et al. (Kitazono et al., 2023). This algorithm hierarchically decomposes a network into the cores with the strongest, second strongest, and third strongest bidirectional connections, and so on. In this figure, the subnetwork in yellow represents the core with the strongest bidirectional connections, whereas that in blue represents the second-strongest core. The entire network shown in purple is the weakest core and has parts with completely unidirectional connections.

Below, we outline the definition of cores according to the method of ref. (Kitazono et al., 2023). A core is defined as a subnetwork that cannot be separated because of its strong bidirectional connections. To define a core, we first introduce the definition of the strength of bidirectional connections. Based on the definition of the strength, we introduce a minimum cut that quantifies the inseparability of a network, which is then used to define cores. Subsequently, we introduce “coreness”—an index, which is defined for each node, that quantifies the strength of the core in which each node is included.

### Strength of bidirectional connections

In this subsection, we introduce a measure that quantifies the strength of bidirectional connections between two parts of a network. Let us consider a directed graph *G*(*V, E*) where *V* denotes the node set and *E* the edge set. For a bi-partition of (*V*_L_*, V*_R_) of the node set *V*, there are two types of edges connecting *V*_L_ and *V*_R_, determined by their direction. One is the set of edges that outgoing from *V*_L_ to *V*_R_:

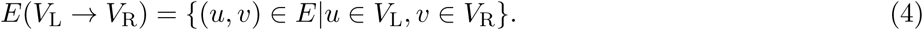

The other is in the opposite direction:

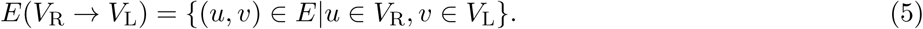

To evaluate the strength of bidirectional connections between *V*_L_ and *V*_R_, we first sum up the weight of the edges in each direction,

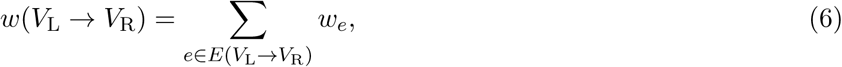

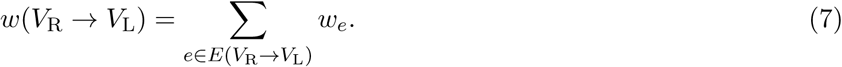

We then define the strength of bidirectional connections as the minimum of the two:

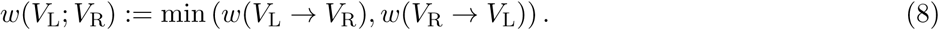

With this definition, when two segments of a network are linked only unidirectionally, the strength of the bidirectional connections *w*(*V*_L_; *V*_R_) equals zero (Figure **1c****(i)**; this indicates that the segments are bidirectionally “disconnected.” In Figure **1c****(ii)**, the connection from one segment to the other is strong (*w*(*V*_L_ → *V*_R_) = 3), whereas the connection in the opposite direction is weak (*w*(*V*_R_ → *V*_L_) = 1). Hence, the bidirectional connections are weak (*w*(*V*_L_; *V*_R_) = 1). In Figure **1c** **(iii)**, the connections in both directions are strong (*w*(*V*_L_ → *V*_R_) = *w*(*V*_R_ → *V*_L_) = 3), resulting in strong bidirectional connections (*w*(*V*_L_; *V*_R_) = 3).

### Measuring inseparability by minimum cut

Using the strength of the bidirectional connections between the two parts of a network defined above, we define a minimum cut (min-cut). As mentioned earlier, a core is a subnetwork that cannot be separated because of its strong bidirectional connections. In other words, a core cannot be cut into two parts without losing many strong edges regardless of how it is cut. To measure such inseparability, we consider the bipartitioning of a network for which the strength of bidirectional connections is the minimum among those for all possible bipartitions—the min-cut. Mathematically, a min-cut 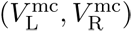 is defined as

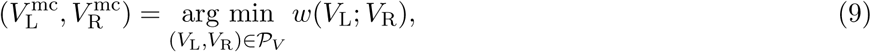

where 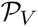 denotes the set of all bipartitions of *V*. We refer to the strength of bidirectional connections for a min-cut as “min-cut weight.” We denote the min-cut weight of a min-cut 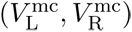 of graph *G* as

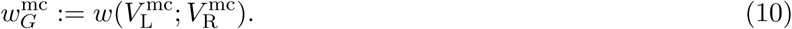

Based on the definition of a min-cut weight, because two parts of a network *G* given by any bipartition are connected with a strength greater than or equal to its min-cut weight 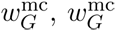 can be taken to represent the network’s inseparability.

### Definition of cores: Complexes

Finally, we introduce the definition of a core (Kitazono et al., 2020; Kitazono et al., 2023). To formally define the cores, we must introduce the concept of an induced subgraph. Let *G* be a graph consisting of a node set *V* and an edge set *E* and let *S* ⊆ *V* be a subset of nodes. Then, an induced subgraph *G*[*S*] is a graph that consists of all the nodes in *S* and all the edges that connect the nodes in *S*. The weight of the minimum cut of *G*[*S*] is denoted by 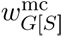. We are now ready to define the complexes as follows.

**Definition 1** (Complex). An induced subgraph G[*S*] (*S* ⊆ *V*) *is called a complex if it satisfies* 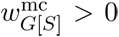 *and* 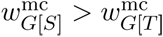 *for any subset T that is a superset of S (T* ⊃ *S and T* ⊆ *V)*.

The definition of complexes is shown schematically in Fig. **1d**, wherein we consider the induced subgraphs of graph *G* that comprises ten nodes {*A, B,…, J*}. An induced subgraph *G*[{*E, F, I, J*}] is a complex because its *w*^mc^ is greater than that of any induced subgraph of *G* that is its supergraph (e.g., *G*[{*B, E, F, I, J*}] and *G*[{*D, E, F, H, I, J*}]). The whole graph *G* is a complex if it satisfies 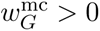 by definition. We define *w*^mc^ = 0 for single nodes because we cannot consider partitions of a single node. Therefore, single nodes cannot be complexes. If we identify complexes by exhaustive search, the computational time increases exponentially with the number of nodes. However, the algorithm proposed by Kitazono et al. (Kitazono et al., 2023) reduces the time complexity of the search to polynomial order (Kitazono et al., 2018; Kitazono et al., 2020; Kitazono et al., 2023), which allows complexes to be extracted even from large networks.

### Coreness

To quantify the strength of the bidirectional connections of the cores wherein each node is included, we measured the “coreness” of each node that we defined using the complexes and their *w*^mc^. A node that is included in complexes with a high *w*^mc^ has high coreness; conversely, a node that is included only in complexes with low *w*^mc^ has low coreness. Specifically, we defined the coreness of a node *v* as *k_v_* if the node *v* is included in a complex with *w*^mc^ = *k_v_* but not included in any complex with *w*^mc^ *> k_v_*. Equivalently, we can define the coreness of a node *v* as the largest of the *w*^mc^ of all complexes containing the node *v*:

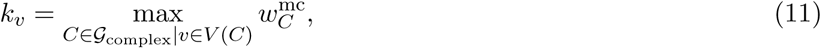

where 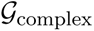 denotes the set of all complexes in the graph *G* and *V* (*C*) denotes the set of all nodes in the complex *C*. The coreness Eq. (11) is equal to the largest of the *w*^mc^ of all subnetworks containing node *v* (Kitazono et al., 2023)

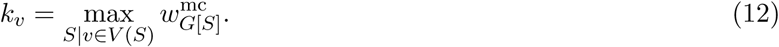

Herein, to investigate the common tendencies of cores across the conditions of the resting state and seven tasks, we normalized coreness by dividing it by the maximum coreness among all nodes for each brain state; then, we averaged these values across all eight brain states. Here, we treated the average of the coreness of Rest 1 and that of Rest 2 as the ‘coreness for Rest.’ It should be noted that the coreness for Rest 1 and Rest 2 were consistent (Extended Data Figure **2-2a**). In the remainder of this paper, “coreness” refers to this averaged value across the conditions.

### Unique characteristics of our core extraction method

Our core extraction method has three essential differences —bidirectionality, globality, and exactness—when compared to a wide range of existing core extraction methods.

#### Bidirectionality

The first difference is that our method is specifically designed to extract cores that are densely bidirectionally connected. To the best of our knowledge, no previous studies have extracted network cores with dense bidirectional connections from brain networks, except for our own study, which analyzed the structural connectome of the mouse brain (Kitazono et al., 2023). We will illustrate how this feature of our method can lead to the differences in extracted cores, particularly by comparing it with s-core decomposition, a representative core extraction method that generalizes the widely used k-core decomposition. To do this, consider the network *W* shown in Extended Data Fig. **1-1a,b**, which is the same as the one shown in Fig. **1d**. In this case, our method extracts only the bidirectionally connected subnetwork (a node set {EFIJ}) as the most central core (Extended Data Fig. **1-1a**), whereas s-core decomposition extracts a network (a node set {CDEFGHIJ}) that includes unidirectionally connected nodes (e.g., nodes C, D, G, H) as the most central core (Extended Data Fig. **1-1b**).

#### Globality

The second distinction is that our method extracts cores by considering the global structure of a network, taking into account whether nodes are interconnected across the entire network, whereas many existing approaches extract cores based on local metrics and cannot account for such network-wide connectivity. Existing core extraction methods, such as degree centrality, network hubs, clustering coefficients, k-core/s-core decomposition explore cores based on local metrics. Specifically, degree centrality, network hubs, and k-core/score decomposition assess cores by focusing on the degree of individual nodes, while clustering coefficients evaluate the connectivity within the local neighborhoods of nodes. Although these approaches effectively capture local structural properties, they do not consider the global structure of the entire network. In contrast, our method identifies cores by taking into account the global structure of the entire network. One example illustrating this difference is the case where a network consists of two modules (Extended Data Fig. **1-1c, d**). There are two modules, each consisting of three nodes, A, B, C, and D, E, F. The modules are connected via one node in each module. In this setting, when considering the global structure, it is natural to expect that two cores ABC and DEF would be extracted. Indeed, in such cases, our method identifies the two sub-networks as the most central cores (Extended Data Fig. **1-1c**). On the other hand, other existing core extraction methods such as k-core/s-core decomposition would identify nearly the entire network as the most central core (Extended Data Fig. **1-1d**).

#### Exactness

The final distinct feature of our method is the exactness of core extraction, whereas other methods are based on approximation. In general, when core extraction methods need to solve a combinatorial optimization problem to find the best set of nodes, the computational time grows exponentially, making it infeasible to explore cores in a realistic amount of time. Therefore, for other core extraction methods that solve such optimization problems, it is generally necessary to approximate core extraction particularly in the case of large network sizes (e.g., modularity maximization (Newman and Girvan, 2004; Newman, 2006; Newman, 2012), participation coefficients (Guimerà and Nunes Amaral, 2005) (which depend on the modularity-based partition), the clique percolation method (Palla et al., 2005; Derényi et al., 2005), or the functional rich club (Deco et al., 2021). For example, performing an exhaustive search for modularity maximization becomes computationally intractable as network size increases, necessitating the use of heuristic or approximate methods, such as the Louvain method (Blondel et al., 2008). By contrast, although our method also solves optimization problems in the process of core extraction, it achieves fast and exact core extraction even from large networks by leveraging an algorithm grounded in the mathematical properties of submodularity and monotonicity (Kitazono et al., 2023).

### Cortical and subcortical rendering

For visualization, the coreness of each ROI was assigned to the cortical surface and the subcortical volume map as follows. First, the coreness of the cerebral cortex was assigned to the fsLR-32k CIFTI space using the parcellation label defined by ref. (Schaefer et al., 2018). The resulting map was then displayed on the “fsaverage” inflated cortical surface by Connectome Workbench (Marcus et al., 2011). For the subcortex, the coreness was assigned to the MNI152 nonlinear 6th-generation space by using the parcellation label from ref. (Tian et al., 2020). The resulting map was then plotted using nilearn (https://nilearn.github.io/).

### Cortical and subcortical major divisions

To analyze the variability in the coreness of ROIs in the cerebral cortex from a cognitive functional perspective, we used the brain atlas developed by Yeo et al. (Yeo et al., 2011), which divides a functional brain network within the cerebral cortex into seven major subnetworks. Each ROI was assigned to one of the seven subnetworks (Schaefer et al., 2018), specifically as: default mode, control, limbic, salience/ventral attention, dorsal attention, somatomotor, and visual networks.

Similarly, the subcortical ROIs were classified according to the Melbourne subcortical atlas (Tian et al., 2020). Each ROI was assigned to one of the seven areas: the hippocampus, amygdala, thalamus, nucleus accumbens, globus pallidus, putamen, and caudate nucleus.

### Comparison of coreness and mean response rates for intracranial electrical stimulation

In this study, we used the mean response rate (MRR) for intracranial electrical stimulation (iES), which is obtained in ref (Fox et al., 2020). The data of ref (Fox et al., 2020) were approved by the Stanford University Institutional Review Board, and the subjects consisted of 67 patients (28 female, mean age ± s.d. = 35.4 ± 12.7 years) selected from a pool of 119 patients admitted to Stanford Hospital for intracranial EEG (iEEG) monitoring of medically refractory epilepsy between 2008 and 2018. Patients were only excluded based on the purely practical considerations, such as the lack of iES sessions, high-quality computed tomography scans, or adequate electrodes covering the cortical gray matter.

In the study of ref (Fox et al., 2020), patients were implanted with either subdural grid/strip electrode arrays (n=53), depth electrodes (stereo-EEG; n=11), or a combination of both (n=3), resulting in a total of 1,476 subdural electrodes and 61 depth electrodes. To ensure precise electrode localization, postoperative CT scans were aligned with preoperative MRI scans and electrodes were linearly projected onto the cortical surface. The standard intrinsic brain network maps (Yeo et al., 2011) encompass only the cortical surface; thus, their analysis incorporated only cortical surface data, excluding subcortical regions. Additionally, although the hippocampus and insula are considered cortical structures, they were excluded due to the complexity of transformation and the potential for seizure induction in these areas.

The mean response rate for iES was obtained as follows: Electrical stimulation was administered to each stimulation site via electrodes. Subjects were then asked to report whether they felt any change in their perceptions, including tactile, visual, emotional perceptions, or motor movement. Electrodes were classified as either “responsive” or “silent” based on the subject’s feedback. Data from 119 subjects were aggregated, and the MRR was calculated for each of the subnetworks of Yeo-17 (or −7) network atlas (Yeo et al., 2011) as the proportion of responsive electrodes compared to all electrodes within each subnetwork. We rendered the MRR for the 17 subnetworks (Fox et al., 2020) on the brain surface in Fig. **3a** and that for the seven networks in Extended Data Fig. **5-1a**. For a comprehensive explanation of the methodology, refer to ref. (Fox et al., 2020).

It is important to note that the term ‘perceptual awareness’ in this study refers to the cognitive aspect of whether perceptual changes induced by iES are recognized, and the scope of this ‘perceptual awareness’ is limited to the types of perceptions elicited. Specifically, the perceptions induced by iES are classified into eight types, as follows: (1) somatomotor effects, (2) visual effects, (3) olfactory effects, (4) vestibular effects, (5) emotional effects, (6) language effects, (7) memory recall, and (8) physiological and interoceptive effects (for more details, please see ref. (Fox et al., 2020)).

To compare coreness with the MRR (Fig. **3c** and Extended Data Fig. **3-1c**), we first calculated the average coreness of the ROIs within each subnetwork of the Yeo-17 network atlas, according to the ROI assignment to each subnetwork by Schaefer et al. (Schaefer et al., 2018), and compared the two metrics. However, prior to this comparison, a minor adjustment to the MRR was necessary owing to the slight differences in voxel allocation to subnetworks between the original Yeo-17 network atlas and Schaefer’s assignment. We projected the MRR onto the brain surface voxels as per the original Yeo-17 network atlas and then recalculated the averages according to Schaefer’s assignment (for details on the calculation method for statistical significance of correlations, see the “Statistical Analysis” section).

### NeuroSynth term-based meta-analysis

To investigate whether the functions of ROIs with high and low coreness in the cerebral cortex differ, we performed a meta-analysis using NeuroSynth—a platform for large-scale automated meta-analysis of fMRI data (www.neurosynth.org) (Yarkoni et al., 2011). We investigated how the degree of association between ROIs and topics that represent cognitive functions vary depending on their coreness as follows. First, we divided the 200 cerebral cortical ROIs into 20 groups according to coreness intervals of 0.05 (0–0.05 to 0.95–1) and, based on which interval respective ROIs belonged to, we classified them into one of these 20 divisions.

Next, for ROIs in each interval, we calculated the association with specific terms as follows (Yarkoni et al., 2011). We first calculated the average value of the “association test” meta-analytic maps provided by Neurosynth (Yarkoni et al., 2011) for the ROIs in each interval. These “association test” maps comprise z-scores obtained from a two-way ANOVA that tests for non-zero associations between voxel activation and the use of a specific term in an article. For example, a large positive z-score for a voxel *i* in the association test map for the term “reward” implies that compared to studies without the term, those with the term in the title or abstract are more likely to report activation of the voxel *i*. By calculating the average of these z-scores for all voxels of ROIs in each interval for every term, we obtain a vector *X* with the dimension of the number of terms that represents the association between the ROIs in the interval and the terms. The voxels were assigned to ROIs by the NIFTI format parcellation label defined in the FSL MNI152 2mm space by Schaefer et al. (Schaefer et al., 2018).

Finally, we evaluated the association of the ROIs in each interval with a specific topic by calculating the Pearson correlation *r* between this vector *X* and a vector *Y*, which is a binary vector assigned for each topic with the dimension of the number of terms, and indicates whether each topic involves the terms. The list of topics and terms, and which term each topic involves, were based on the data available at https://github.com/NeuroanatomyAndConnectivity/gradient_analysis/blob/master/gradient_data/neurosynth/v3-topics-50-keys.txt. The number of topics was 50, of which we used 44 topics for analysis, after excluding six topics that did not capture any consistent cognitive functions following ref. (Margulies et al., 2016). The topic terms (e.g., “eye movements,” “cued attention,” and “emotion”) that represent the topics were set according to Margulies et al. (Margulies et al., 2016). Fisher’s z-score, obtained by Fisher’s z-transformation of the correlation *r* between the vectors *X* and *Y*, was used as the degree of association between each topic and ROIs in the interval. For more information see https://neurosynth.org and ref. (Yarkoni et al., 2011).

We calculated Fisher’s z-score for all intervals and all topics. Subsequently, for visualization, we sorted the topics based on the weighted average of the coreness of intervals by using Fisher’s z-score as the weight, wherein topics related to high- and low-coreness ROIs were placed at the top and bottom, respectively. Any topic that did not reach a significant threshold of *z >* 3.1 in any interval was excluded; thus, only the remaining 24 topics were displayed (Fig. **4**). Therefore, while the range of cognitive functions specified by the term ‘cognition’ in this study is limited to these 24 topics, it should be noted that these topics cover a range from lower-order sensorimotor functions (e.g., motor, eye movement, visual perception, and auditory processing) to higher-order cognitive functions (e.g., social cognition, verbal semantics, and autobiographical memory). The analysis was performed using a modified code available at https://github.com/NeuroanatomyAndConnectivity/gradient_analysis.

### Comparison of the functional connectivity gradient and coreness

A functional connectivity gradient refers to the spatial variation in functional connectivity patterns across different brain regions (Margulies et al., 2016; Müller et al., 2020; Shafiei et al., 2020; Fornito et al., 2019). Mathematically, a functional connectivity gradient is obtained through an embedding of a functional network into a low-dimensional space. Previous studies have shown that functional connectivity gradients are associated with numerous neuroscientific features (Margulies et al., 2016; Müller et al., 2020; Shafiei et al., 2020; Fornito et al., 2019), including molecular, cellular, anatomical, and functional aspects, indicating that these gradients capture essential properties of the brain’s functional organization. Among those functional connectivity gradients, in this study, we utilized the one for the entire human cerebral cortex developed by Margulies et al. (Margulies et al., 2016), which was obtained by the following procedures: A functional connectivity matrix was first obtained by calculating the correlation between all pairs of gray coordinates from resting-state fMRI data. Next, a nonlinear dimensionality-reduction technique called diffusion embedding (Coifman et al., 2005) was applied to this connectivity matrix. The functional connectivity gradient was defined as the first component in the embedding space, which accounts for the greatest variance in the connectivity patterns. Margulies et al. have demonstrated that the functional connectivity gradient corresponds to a cortical spatial gradient of the degree of abstraction and integration in processing, ranging from the primary sensory/motor cortex to transmodal areas (Margulies et al., 2016). The detailed methodology for calculating the functional connectivity gradient has been described in ref. (Margulies et al., 2016). Data of the functional connectivity gradient are publicly available at https://github.com/NeuroanatomyAndConnectivity/gradient_analysis. For our analysis, we averaged the functional connectivity gradient for the voxels in each ROI, and these averaged values were then used for the scatterplot in Fig. **5c** (for details on the calculation method for statistical significance of correlations, see the “Statistical Analysis” subsection).

It should be noted that the sign of the gradient is reversed from that in Margulies et al.’s paper but matches that of the gradient data available at the repository. This difference arises because the sign of the functional connectivity gradient is arbitrary and carries no particular meaning. In diffusion embeddings, each axis’s sign does not convey specific information due to the nature of the eigenvectors used in constructing the embedding. In this context, eigenvectors are determined only up to a sign, meaning that each axis can be flipped independently without altering the structure of the embedding.

### Extraction of cores regardless of bidirectionality

To assess the impact of incorporating the bidirectionality of connections on the results, we extracted cores when bidirectionality was ignored. Instead of using Eq. (8), which defines the measure of bidirectional connection strength, we used a simpler measure: we summed the weights of all edges that connect two parts, regardless of their directions:

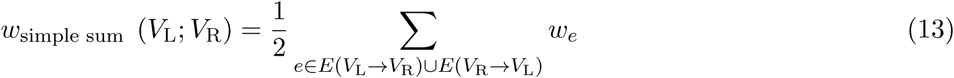

Here, the factor of 2 in the denominator is to maintain consistency with the case wherein bidirectionality is considered. The use of this connection strength Eq. (13) is equivalent to applying the bidirectional connection strength Eq. (8) to an undirected network, which is obtained by ignoring edge directions and is equivalently achieved by setting its connection matrix to *W*^′^ = (*W* + *W*^⊤^)*/*2, where *W* is the connection matrix of the original directed network (Kitazono et al., 2023).

### Weighted degree of nodes

The weighted degree deg(*v*) of a node *v* is defined as the sum of the weights of all edges connected to the node *v*, regardless of the edge directions:

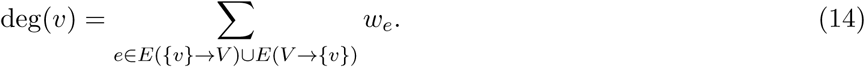

To investigate the common tendencies of node degree across the conditions of the resting state and seven tasks, we normalized weighted degree of nodes in the same way as we normalized coreness: by dividing it by the maximum degree among all nodes for each brain state and then averaging these values across all brain states. In the remainder of this paper, “weighted degree” refers to this averaged value across the conditions.

### Statistical Analysis

#### Comparison of Coreness Between Cortical and Subcortical Regions

We performed a one-way ANOVA to compare the coreness between cortical and subcortical regions to examine the differences between cortical and subcortical networks (Fig. **2b**, Fig. **6b**).

**Figure 2:**
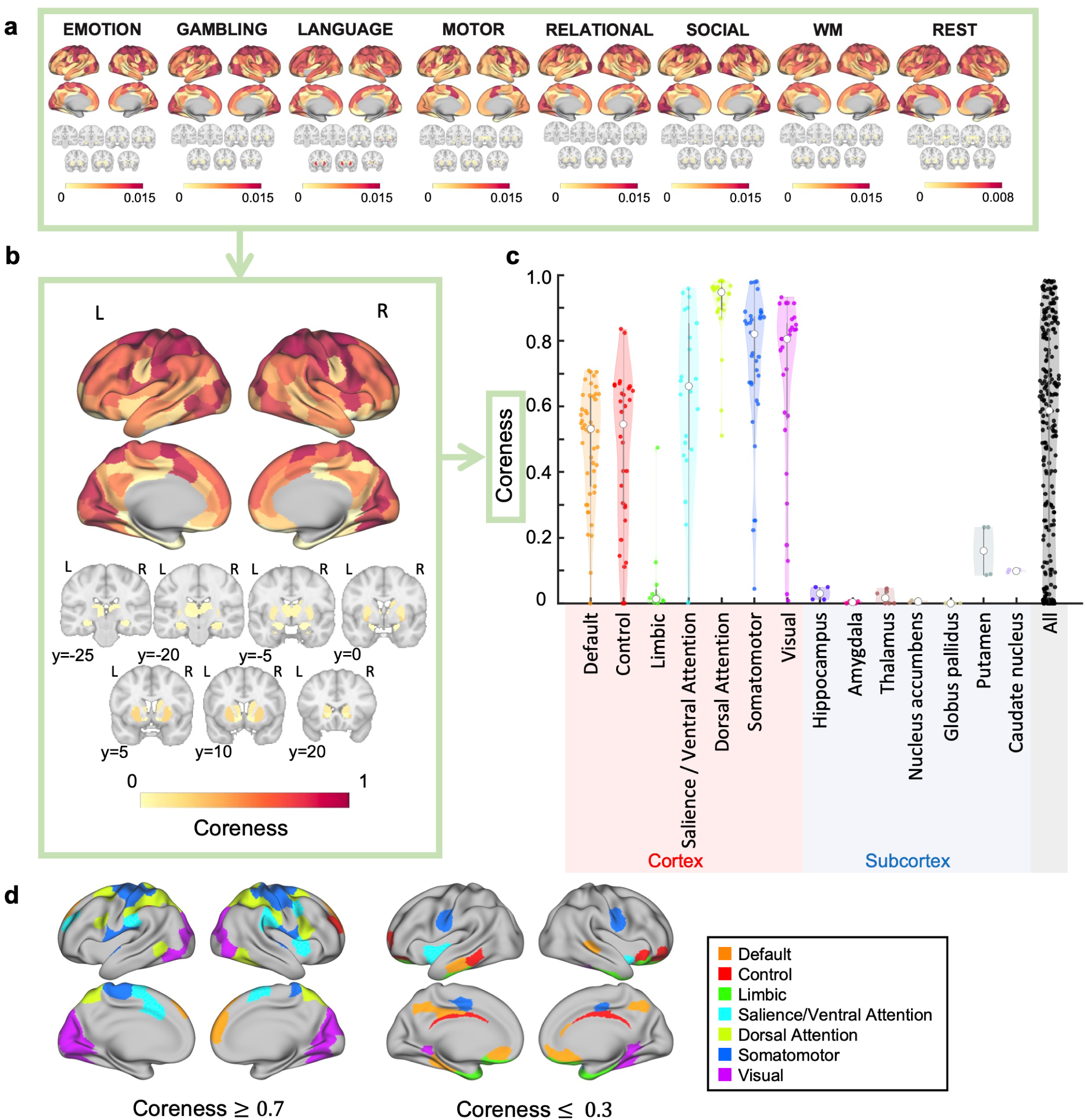
Extracted bidirectionally interacting cores. Compared to subcortical regions, cerebral cortical regions tend to have higher coreness. Furthermore, within the cerebral cortex, ROIs with high coreness are particularly found in the salience/ventral attention, dorsal attention, somatomotor, and visual networks. In contrast, ROIs in the limbic network show low coreness, and the default mode network and frontoparietal control network include ROIs with low coreness. **a**, Coreness at rest and during seven tasks in the cerebral cortex (top) and the subcortex shown in seven coronal slices (bottom). Here, coreness is not normalized. **b**, Coreness in the cerebral cortex (top) and the subcortex shown in seven coronal slices (bottom). Coordinates of the slices were given in the MNI (Montreal Neurological Institute) space. **c**, Violin plots of coreness according to the divisions of the Yeo-7 network atlas and major divisions of the subcortex. Each violin plot represents the probability density of coreness for each division, and the colored dots inside the plots represent ROIs. The white dot represents the median, and the thick line inside indicates the interquartile range. **d**, ROIs with coreness *>* 0.7 or *<* 0.3 are colored as per the Yeo-7 network atlas.

#### Principles of statistical analysis in comparing coreness with other metrics

Given that our comparisons of coreness with the functional gradient and the MRR for iES are exploratory rather than hypothesis-driven, and considering the recommendation against interpreting results solely based on p-values (Wasserstein and Lazar, 2016), we avoid a dichotomous judgment of the significance of results based on p-values alone. Instead, we report all relevant information, such as 95% confidence intervals and central 95% interval of the null distribution based on brainSMASH to provide readers with a more comprehensive and accurate interpretation of the results. Similarly, for regression analyses, we provide 95% confidence intervals. The details of the statistical analyses are described in the following paragraphs.

#### Calculation of confidence intervals for correlations

To calculate the 95% CIs, we used the bias-corrected and accelerated percentile (BCa) bootstrap method (Efron and Tibshirani, 1994), with 10,000 bootstrap samples.

#### Calculation of p-values for Correlation Analysis

Simple statistical analyses for the Pearson correlations may result in false positives due to spatial autocorrelation in brain maps. Therefore, we used brainSMASH toolbox (Burt et al., 2020) to generate null maps that consider spatial autocorrelation in cortical regions, in order to test the significance of the correlations between the coreness map and either the iES map or Margulies maps. We generated 10,000 surrogate maps and constructed the null distribution of correlations by calculating the correlations between the empirical data (the mean response rate for iES or the functional connectivity gradient) and these surrogate maps. From this null distribution, we defined the central 95% interval of the null distribution as the range between the 2.5% and 97.5% percentiles. For the comparison with the iES map, we averaged the surrogate map values within each partition of the 17 Yeo network (or 7 Yeo network) and obtained the null distribution of correlation values for the surrogate data map. Then, we calculated p-values (referred to as *p*_brainSMASH_) based on where the empirical correlation value fell within the null distribution.

#### Calculation of the confidence intervals for the regression line

In the scatter plots of coreness with either mean response rates for iES or functional connectivity gradient, the 95% confidence interval for the regression line was calculated using the following formula: 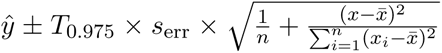 where *ŷ* is the predicted value on the regression line, *T*_0.975_ is the 97.5th percentile of the Student’s t-distribution with *n* − 2 degrees of freedom, *s*_err_ is the standard error of the residuals, *n* is the sample size, *x* is the explanatory variable, and *x̄* is the mean of *x*.

## Data and code availability

The neuroimaging data are freely available from HCP https://db.humanconnectome.org/. Parcellation labels were used from https://github.com/ThomasYeoLab/CBIG (for the cerebral cortex) and https://github.com/yetianmed/subcortex (for the subcortex). The MATLAB codes for extracting bidirectionally connected cores are available at https://github.com/JunKitazono/BidirectionallyConnectedCores. The estimation of the VAR model was performed using the MVGC toolbox (Version 1.2) https://github.com/SacklerCentre/MVGC1. Normalized directed transfer entropy (NDTE) was calculated using a modified version of the codes at https://github.com/gustavodeco/nhb-ndte. For the visualization of regions within the cerebral cortex and subcortex, the connectome workbench https://www.humanconnectome.org/software/connectome-workbench and nilearn https://github.com/nilearn/nilearn, respectively, were used. For the neurosynth meta-analysis, we used a modified version of the codes at https://github.com/NeuroanatomyAndConnectivity/gradient_analysis. Violin plots were drawn using the codes at https://github.com/bastibe/Violinplot-Matlab.

## Results

### Framework for extracting bidirectionally interacting cores from a directed functional network

This section outlines a novel framework that has been proposed in this paper for extracting the cores of brain networks, with strong bidirectional interactions from brain activity data. The framework consists of two steps. First, the strength of statistical causal influence between brain regions is estimated from brain activity to construct a whole-brain-directed functional network. Second, strongly bidirectional cores are extracted from the network.

In the first step, we quantify the strength of statistical causal influence from one ROI to another using fMRI data (Fig. **1a**). This quantification is performed for every possible combination of ROIs, enabling us to construct a whole-brain directed network (Figure **1b**) that illustrates the strength of statistical causal influences among brain ROIs. To measure these strengths, we used normalized directed transfer entropy (NDTE) (Deco et al., 2021)—a statistical method that is designed to estimate the strengths of the statistical causal influence between two sets of time-series data. The direction of the arrow in the schematic (Figure **1b**) indicates the direction of statistical causal influence between ROIs, whereas the thickness of these arrows reflects the magnitude of the influence. The methodology for this calculation has been described in the Methods section.

In the second step, from the constructed directed network, we extract subnetworks with strong bidirectional interactions, or the “cores” of a network that are identified using the method proposed by Kitazono et al. (Kitazono et al., 2023). In their study, a core is termed a “complex” (Kitazono et al., 2023), which is defined as a subnetwork that is composed of stronger bidirectional connections than other subnetworks that include it. Using the method, the directed network can be hierarchically decomposed into complexes based on the strength of bidirectional connections. Here, the strength of bidirectional connections is defined based on a measure that quantifies how strongly the 2 divided parts of a network are bidirectionally connected, as shown in Fig. **1c** (See Methods for details). The complexes extracted from the network in Fig. **1b** are shown in Fig. **1e**, and an example using a toy network is shown in Fig. **1d** (See Methods for details).The subnetwork highlighted with a yellow background is the complex with the strongest bidirectional connections; the subnetworks distinguished by blue and purple backgrounds represent the second-strongest and the weakest complexes, respectively. Generally, complexes exhibit a nested structure wherein stronger bidirectional complexes are contained in weaker ones to form either unimodal or multimodal hierarchical structures.

To quantify the strength of the bidirectional connections of the cores wherein each node is included, we use a measure called the “coreness.” We defined the coreness of a node *v* as the largest of the *w*^mc^ of all complexes containing the node *v* (see Methods for details).

### Bidirectional cores in the functional network of the human brain

In this subsection, we present the results of our analysis of fMRI data from the human connectome project (HCP) (Van Essen et al., 2013) using our proposed framework. We assume that if a brain region is consistently included in the strongly bidirectional, central cores during the execution of various tasks, those regions are crucial for diverse cognitive functions in general. Therefore, to determine which regions are consistently included in central cores, we first extracted cores at rest and during seven tasks in the HCP. Next, to explore how regions consistently included in the central cores relate to perceptual awareness, we compared coreness with the rates of iES-induced perceptual elicitation. Additionally, to further characterize the regions consistently included in the central cores, we performed a meta-analysis using NeuroSynth, and compared the cores with the functional connectivity gradient.

### Brain regions frequently included in the central cores

We applied our proposed framework to HCP fMRI data to analyze which brain regions were more likely, on average, to be included in the central cores. First, we extracted cores for the resting state and seven tasks (Figure **2a**, Extended Data Fig. **2-1**). Subsequently, we calculated the average coreness over these eight conditions (see Methods for details). This average is simply referred to as coreness, hereafter. It should be noted that we treated the average of the coreness of Rest 1 and that of Rest 2 as the coreness for resting state, after confirming their consistency (Extended Data Figure **2-2a**).

The results revealed that compared to the subcortical regions, cerebral cortical regions tend to have higher coreness (*F* (1, 230) = 115.295*, p* = 4.59 × 10^−22^*, η*^2^ = 0.334; Figs. **2b** and **2c**). Many ROIs in the cerebral cortex display high coreness, whereas all subcortical regions show lower coreness, although they are from diverse areas, such as the hippocampus, amygdala, and thalamus. This trend remained broadly consistent even when the graph density was varied to 5% and 20% (one-way ANOVA, 5%; *F* (1, 230) = 71.187, *p* = 3.64 × 10^−15^, *η*^2^ = 0.236, 20%; *F* (1, 230) = 170.032*, p* = 1.83 × 10^−29^*, η*^2^ = 0.425, Extended Data Figs. **2-3a** and **b**), indicating the robustness of the findings across different thresholds. This tendency indicates that although bidirectional cores are formed within cortical regions, subcortical regions do not directly become part of these cores.

A more detailed examination within the cerebral cortex revealed that not all ROIs possess high coreness, indicating variability in coreness among different regions (Fig. **2b**). To understand this variability in coreness from a cognitive functional perspective, we divided ROIs into seven functional subnetworks according to Yeo’s 7 network atlas (Yeo et al., 2011) (Fig. **2c**). The results showed that ROIs with high coreness were particularly located in the salience/ventral attention, dorsal attention, somatomotor, and visual networks. Conversely, ROIs in the limbic network exhibited low coreness, and the default mode network and control network included ROIs with low coreness.

Next, to explore the spatial distribution trend of ROIs with particularly high or low coreness within the cerebral cortex, we visualized ROIs with a coreness greater than 0.7 and less than 0.3 on the surface of the cerebral cortex (Fig. **2d**). This visualization used color coding based on the Yeo-7 network atlas. The results showed that ROIs with high coreness were mainly located in the somatosensory and motor areas surrounding the central sulcus, which are part of the somatomotor and dorsal attention networks, and in the occipital areas, which belong to the visual network. Additionally, high-coreness ROIs were identified in the salience network, which were relatively dispersed in their locations. On the other hand, low-coreness ROIs were scattered across various areas, including the lateral temporal, medial temporal, orbitofrontal, and cingulate cortices. This trend was confirmed to remain fundamentally unchanged even when the coreness thresholds were changed (coreness *>* 0.8 and *<* 0.2, or *>* 0.6 and *<* 0.4; Extended Data Fig. **2-3c**).

### Comparison of the core structure with the mean response rate for iES

Next, we investigated whether the coreness of cerebral cortical ROIs, whether high or low, is associated with their importance for perceptual awareness. Specifically, we compared the coreness with the rates of iES-induced perceptual elicitation (Fox et al., 2020).

Figure **3a** shows the cortical map representing the mean response rate (MRR) for iES. By comparing this map of the MRR with the map of coreness (Fig. **3b**, the same figure as Fig. **2b** is reprinted), similar structures can be observed in the two maps—high MRR and coreness are commonly found in the areas such as the somatosensory and motor areas surrounding the central sulcus, and in the visual cortex in the occipital region, whereas low MRR and coreness are visible in the areas such as the prefrontal, lateral temporal, posterior parietal, and posterior cingulate cortices. The scatterplot (Fig. **3c**) shows an apparent upward trend between coreness and MRR, resulting in a moderate positive correlation *r* = 0.462. Note, however, that the correlation estimate is not statistically robust due to the small sample size (*n* = 17). The 95% confidence interval (CI) and the central 95% interval of the null distribution based on brainSMASH are very wide, [-0.074, 0.755] and [-0.583, 0.574], respectively, and the corresponding p-value computed from the null distribution is *p*_brainSMASH_ = 0.0863. A positive correlation was observed not only in the comparison using the Yeo-17 network atlas (Fig. **3c**) but also in the comparison using the Yeo-7 network atlas (*r* = 0.567), although the uncertainty of the correlation estimate is even larger because of the smaller sample size (95% CI [-0.994, 0.944], 95% null distribution interval (brainSMASH) [-0.800, 0.796], *p*_brainSMASH_ = 0.139, *n* = 7 points, Extended Data Figs. **3-1a, b, c**).

**Figure 3:**
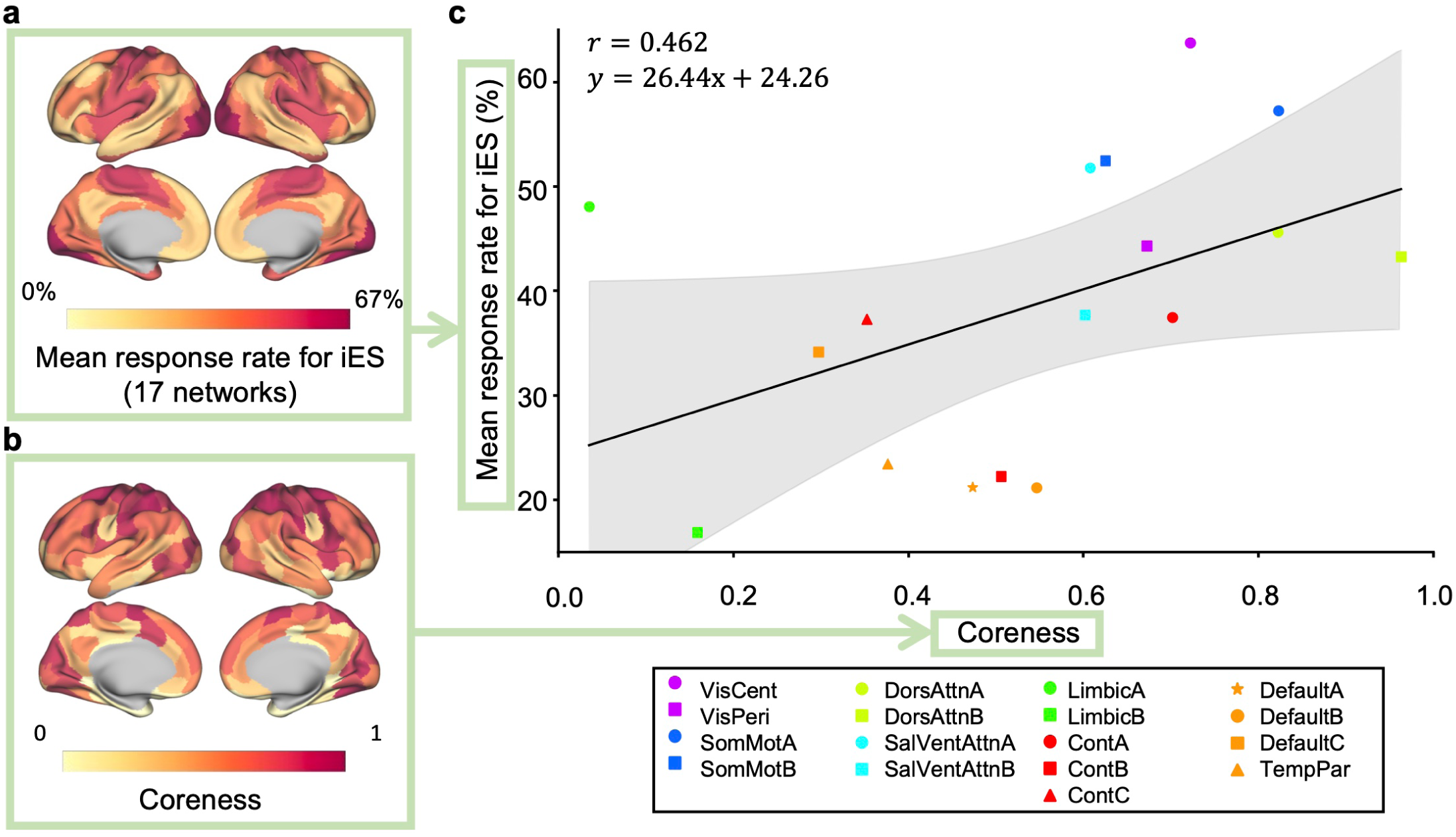
Comparison of the core structure with the mean response rate for iES. There is a correlation between coreness and the mean response rate (MRR) for iES. **a, b**, A cortical surface rendering of the MRR for iES (Fox et al., 2020) (**a**) and that of coreness (**b**, same as Fig. **2b**). **c**, A scatter plot showing the average coreness for each division of the Yeo-17 network atlas (horizontal axis) and the MRR (vertical axis). A positive correlation exists between the coreness and the MRR (*r* = 0.462). The solid line in the figure represents the least-squares line and the shaded area represents the 95% confidence interval. Each point is color-coded according to the Yeo-7 network atlas and then assigned marker shapes according to the Yeo-17 network atlas.

The moderate level of positive correlation between coreness and MRR was also observed even when the graph density was changed to 5% and 20%, showing the robustness of the observed trend with resepect to the graph density (5% graph density: *r* = 0.498, 95% CI [0.043, 0.767], 95% null distribution interval (brainSMASH) [-0.583, 0.574], *p*_brainSMASH_ = 0.0623, *n* = 17 points; 20% graph density: *r* = 0.384, 95% CI [-0.176, 0.717], 95% null distribution interval (brainSMASH) [-0.544, 0.600], *p*_brainSMASH_ = 0.165, *n* = 17 points; Extended Data Figs. **3-1d** and **3-1e**).

### Term-based meta-analysis of the core structure using NeuroSynth

To further explore the cognitive functions linked to ROIs with varying levels of coreness, we conducted a term-based meta-analysis using NeuroSynth (Yarkoni et al., 2011), which statistically evaluates the association between each ROI and cognitive functions in the literature by analyzing information from thousands of published fMRI-based studies.

As an example, we illustrate the results obtained by applying a Fisher’s z-score threshold of 5, chosen conservatively to account for multiple comparisons (approximately 500 comparisons in the meta-analysis table). Under this criterion, high-coreness ROIs (e.g., coreness ≥ 0.75) are associated with terms such as ‘eye movements,’ ‘numerical cognition,’ ‘visual attention,’ ‘visual perception,’ ‘reading,’ ‘action,’ ‘motor,’ ‘cued attention,’ ‘working memory,’ ‘multisensory processing,’ ‘cognitive control,’ ‘visuospatial,’ ‘pain,’ and ‘auditory processing’ (Fig. **4**). Most of these terms reflect lower-order sensorimotor cognitive functions, although some terms (like ‘reading’, ‘numerical cognition’, ‘working memory’) may rely on both sensorimotor and higher-order components. In contrast, low-coreness ROIs (e.g., coreness ≤ 0.25) are associated with terms such as ‘face/affective processing,’ ‘autobiographical memory,’ ‘emotion,’ ‘declarative memory,’ ‘visual semantics,’ ‘pain,’ and ‘motor,’ based on the same z-score criterion (Fig. **4**). These terms primarily correspond to higher-order cognitive processes, though ‘pain’ and ‘motor’ are more closely tied to lower-order sensorimotor functions. This trend was similarly observed when the graph density was changed to 5% and 20% (Extended Data Figs. **4-1b** and **4-1c**), and when subcortical regions were included (Extended Data Figure **4-1a**).

**Figure 4:**
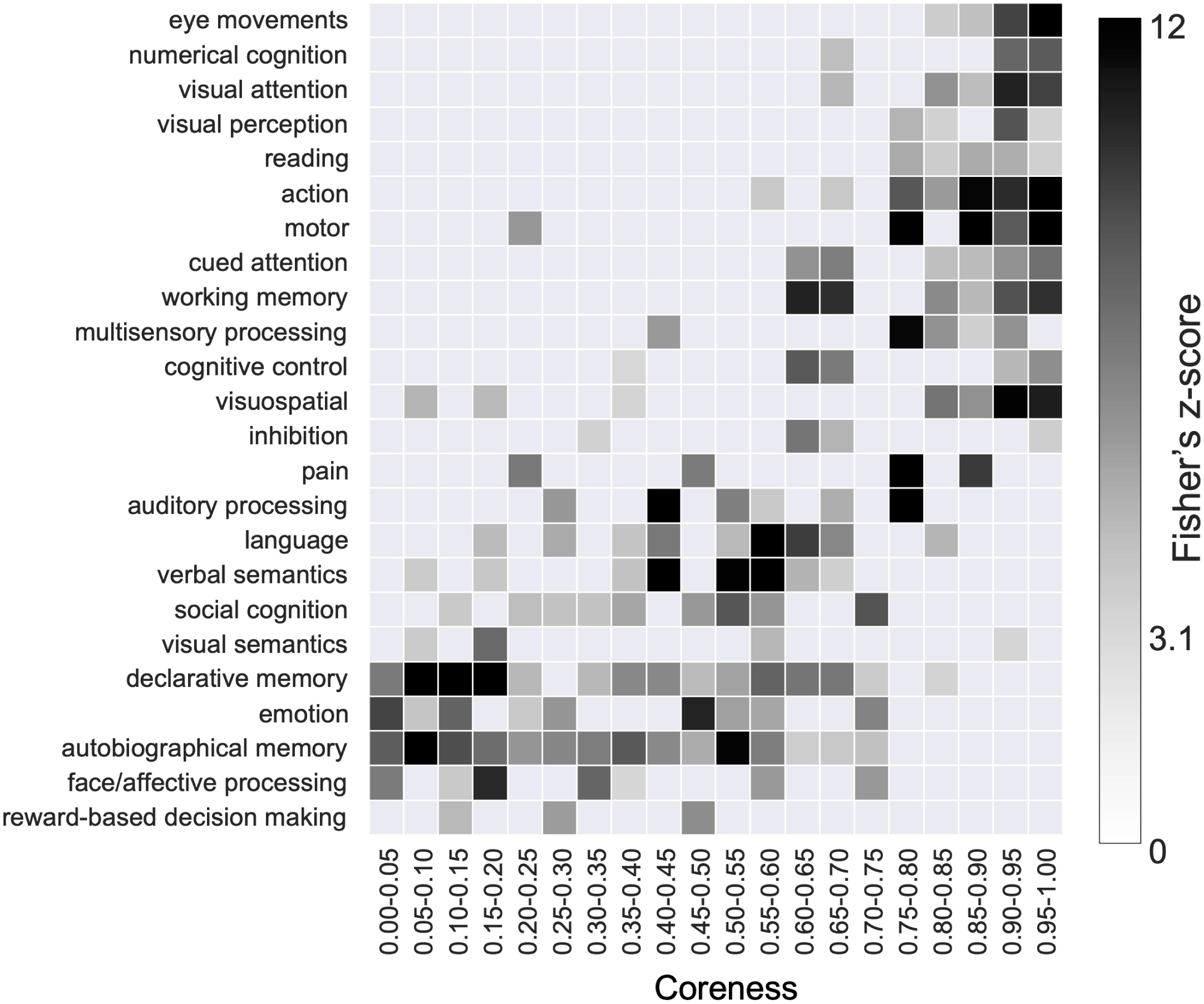
Term-based meta-analysis of the core structure using NeuroSynth. High-coreness ROIs are related to lower-order sensorimotor functions whereas low-coreness ROIs are related to higher-order cognitive functions. This figure shows a term-based meta-analysis using NeuroSynth applied to the coreness of the cerebral cortex. The columns represent coreness, at intervals of 0.05 from 0–0.05 to 0.95–1. The rows represent the topic terms used in the meta-analysis. The grayscale of each cell indicates the Fisher’s z-score representing the association strength between ROIs in each division of coreness and topic terms, as obtained from the meta-analysis. Only components that reached a significant threshold of *z >* 3.1 are colored. For visualization, topic terms are arranged by the weighted mean of the coreness of intervals with Fisher’s z-score as weights, by placing terms related to high-coreness regions at the top and those related to low-coreness regions at the bottom. Only the cerebral cortical ROIs were used for this analysis (see Methods for details).

### Comparison of the core structure with the functional connectivity gradient

To further assess the relationship between coreness and lower-order or higher-order functions in the cerebral cortex, we compared coreness with the functional connectivity gradient (Margulies et al., 2016), which is a low-dimensional embedding of the functional connectivity at rest and accounts for the greatest variance in the connectivity patterns. The gradient is known to correspond to a spectrum of the degree of abstraction and integration in processing, where its upper end is associated with lower-order sensory processing and the lower end with higher-order (abstract and integrative) cognitive functions (Margulies et al., 2016). It should be noted that this is the inverse of the gradient described by Margulies et al. (Margulies et al., 2016).

Therefore, if there exists a relationship where high-coreness and low-coreness ROIs are associated with lower-order and higher-order functions, respectively, we would expect the coreness to correlate with the functional connectivity gradient. This expectation was confirmed by the comparison of the two (see Fig. **5a** and Fig. **5b**, Fig. **2b** reprinted). Regions at one end of the gradient comprise visual, somatosensory/motor, and auditory areas, demonstrating partial overlap with regions of high coreness. Similarly, regions at the other end of the gradient predominantly comprise default-mode network areas, which show partial overlap with low-coreness regions. This relationship is further evidenced in the scatterplot comparing coreness with the functional connectivity gradient (Fig. **5c**), indicating a statistically robust positive correlation between them (*r* = 0.397, 95% CI [0.299, 0.478], the central 95% interval of the null distribution (brainSMASH) [-0.324, 0.311], *p*_brainSMASH_ = 0.00280, *n* = 200 points). The positive correlation was also observed even when the graph density was changed to 5% and 20%, showing the robustness of the observed trend with resepect to the graph density (5% graph density: *r* = 0.441, 95% CI [0.341, 0.527], 95% null distribution interval (brainSMASH)[-0.324, 0.311], *p*_brainSMASH_ = 0.00100, *n* = 200 points; 20% graph density: *r* = 0.304, 95% CI [0.207, 0.386], 95% null distribution interval (brainSMASH) [-0.304, 0.335], *p*_brainSMASH_ = 0.0439, *n* = 200 points; Extended Data Fig. **5-1a** and **5-1b**).

**Figure 5:**
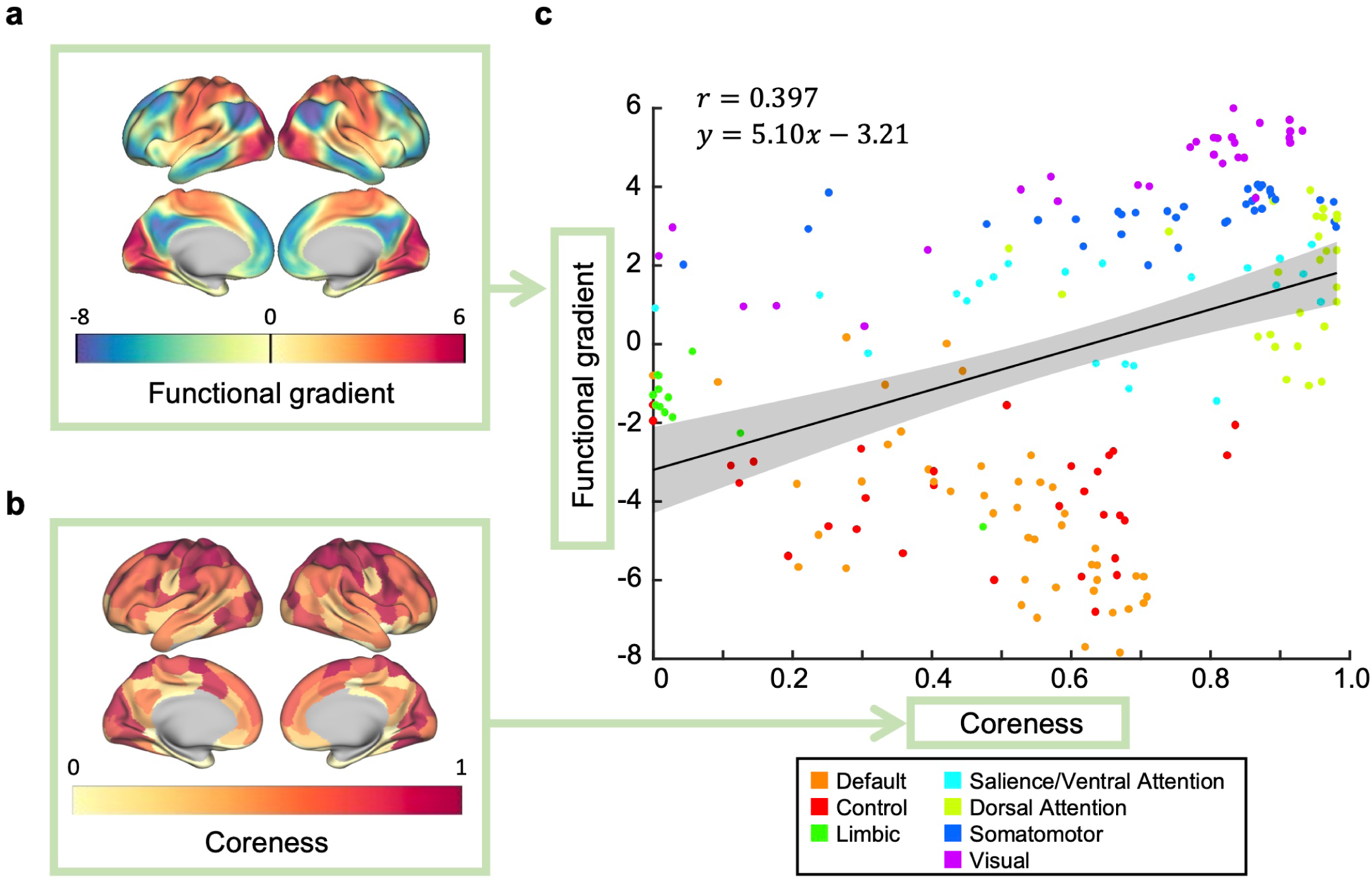
Comparison of the core structure with the functional connectivity gradient. The functional connectivity gradient (Margulies et al., 2016) and coreness are correlated. **a, b**, A cortical surface rendering of the cortical functional connectivity gradient (Margulies et al., 2016) (**a**) and that of coreness (**b**, same as Fig. **2 b**). **c**, A scatterplot of coreness and the functional connectivity gradient. A positive correlation exists between coreness and the functional connectivity gradient (*r* = 0.397). The solid line represents the least-squares line and the shaded area represents the 95% confidence interval. Each point represents an ROI and is color-coded according to the Yeo-7 network atlas (See Methods for details).

### Comparison with the complexes when bidirectionality is ignored

In this subsection, we explore the importance of considering bidirectionality in findings presented in the previous sections. To do so, we examined how cores would change if we only considered the strength of interactions while ignoring bidirectionality. Not accounting for bidirectionality equates to the symmetrizing of a network and treating it as an undirected network (see Methods for details).

First, on comparing the broad categories—cortical and subcortical—similar to the observations made when considering bidirectionality (Figs. **2b** and **2c**), cortical regions generally exhibited higher coreness than subcortical regions, even in the absence of consideration of bidirectionality (*F* (1, 230) = 235.492*, p* = 4.52×10^−37^*, η*^2^ = 0.506; Figs. **6a** and **6b**). Next, we examined the differences within the cerebral cortex when considering, versus ignoring, bidirectionality. The results showed that although coreness was correlated between the two cases, large differences were observed in certain regions (Fig. **6c**); specifically, in regions with low or medium coreness when considering bidirectionality, coreness was largely increased by ignoring bidirectionality (see Fig. **6c**, highlighted area, and Fig. **6d**). Thus, there was a change that counteracted the trend in coreness that existed when bidirectionality was considered. This suggests that the correlation of coreness with the MRR for iES, as well as with the functional connectivity gradient—both observed when considering bidirectionality—weakened when bidirectionality was ignored. Indeed, in regions with smaller MRR and the functional connectivity gradient tended to show a greater increase in coreness if bidirectionality was ignored (MRR; *r* = −0.373, 95% confidence interval (CI) [-0.675, 0.080], the central 95% interval of the null distribution based on brainSMASH [-0.588, 0.552], *p*_brainSMASH_ = 0.148, computed on *n* = 17 points, functional connectivity gradient; *r* = −0.256, 95% CI [-0.377, −0.125], 95% null distribution interval (brainSMASH) [-0.308, 0.278], *p*_brainSMASH_ = 0.0560, *n* = 200 points, Extended Data Figs. **6-1a** and **6-1b**), resulting in a weaker correlation of coreness with the MRR and with the functional connectivity gradient (MRR; *r* = 0.419, 95% CI [-0.139, 0.723], 95% null distribution interval (brainSMASH) [-0.536, 0.593], *p*_brainSMASH_ = 0.138, *n* = 17 points, functional connectivity gradient; *r* = 0.357, 95% CI [0.276, 0.425], 95% null distribution interval (brainSMASH) [-0.296, 0.324], *p*_brainSMASH_ = 0.00990, *n* = 200 points, Extended Data Figs. **6-1c** and **6-1d**).

**Figure 6:**
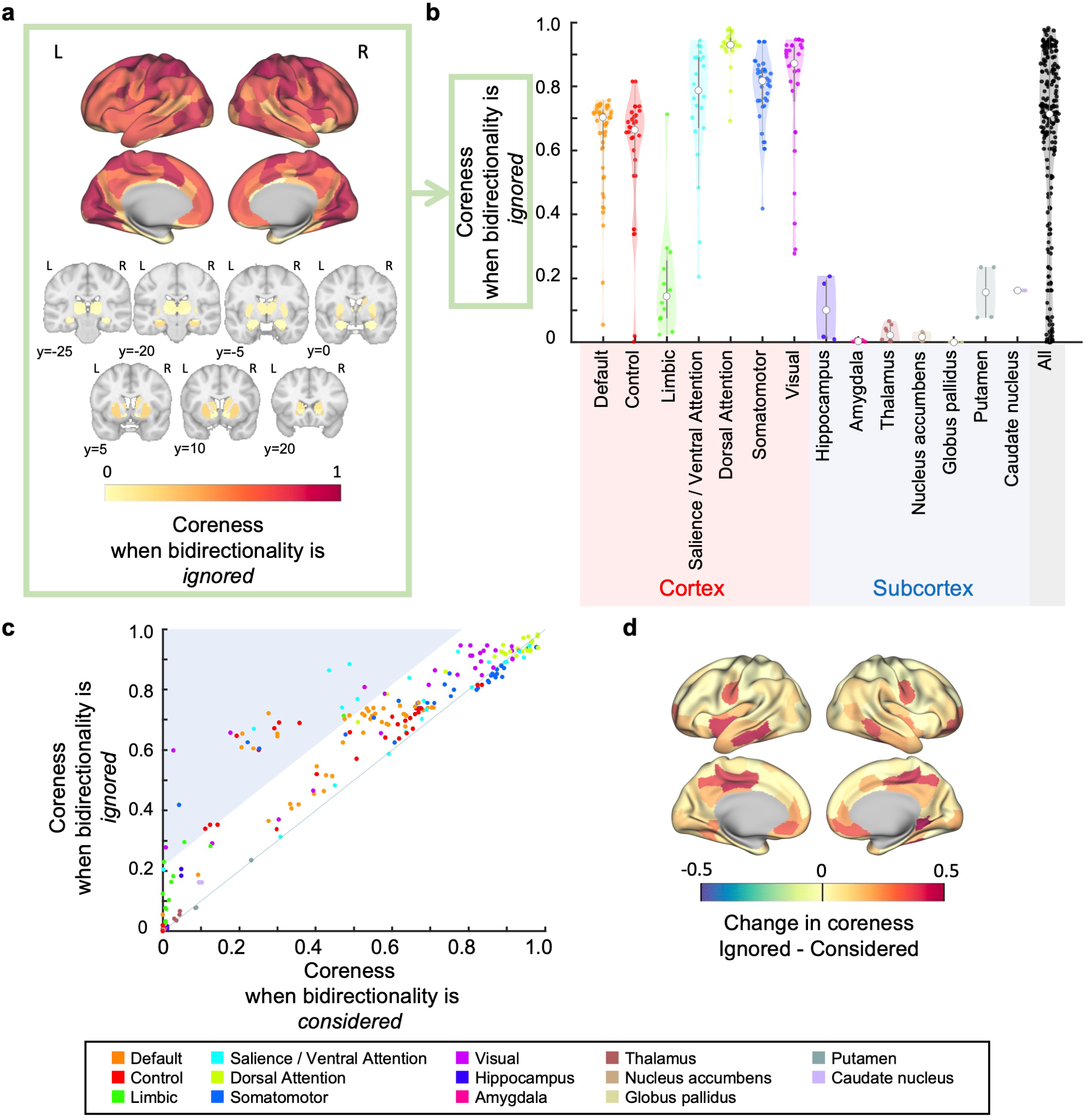
Comparison of the core structure with the case when bidirectionality is ignored. **a** and **b**, When bidirectionality is ignored, the cortical regions tend to have higher coreness compared to the subcortical regions, as in the case when bidirectionality is considered. **a**, A cortical surface rendering of coreness when bidirectionality is ignored in the cerebral cortex (top) and that in the subcortex shown in seven coronal slices (bottom). The coordinates of the slices were given in the MNI space. **b**, Violin plots of coreness when bidirectionality is ignored according to the divisions of the Yeo-7 network atlas and major divisions of the subcortex. **c**, A comparison of coreness between considering and ignoring bidirectionality. The difference between the two cases is small and large for regions with high and low coreness when bidirectionality is considered, respectively. Each point is an ROI and is color-coded according to the Yeo-7 network atlas and subcortical divisions. The solid line represents the identity line (*y* = *x*). The blue background highlights regions with relatively large changes in coreness between considering and ignoring bidirectionality. **d**, A cortical surface rendering of the changes in coreness.

### Comparison with other existing methods for extracting cores

To further assess the importance of considering bidirectionality, we compared our core extraction method using complexes with other existing methods that do not consider bidirectionality. Specifically, we conducted comparisons with *s*-core decomposition (Chatterjee and Sinha, 2007; van den Heuvel and Sporns, 2011; Harriger et al., 2012; Crobe et al., 2016) and network hubs (van den Heuvel and Sporns, 2013; Royer et al., 2022).

First, we undertook a comparison with the *s*-core decomposition. The *s*-core decomposition is frequently used for extracting subnetworks with strong connections, which are called *s*-cores, and does not consider the bidirectionality of connections (Chatterjee and Sinha, 2007; van den Heuvel and Sporns, 2011; Harriger et al., 2012; Crobe et al., 2016). The comparison revealed that the coreness for the complexes when bidirectionality is considered and the coreness for *s*-core decomposition (where coreness can be defined in the same way as the complexes. See Methods for details.) were not necessarily equal (Fig. **7a**). This indicates that, by considering bidirectionality, we can extract core structures that are not identifiable by *s*-core decomposition. Additionally, the coreness for *s*-cores and that for complexes when bidirectionality is ignored were approximately equal (Fig. **7b**).

**Figure 7:**
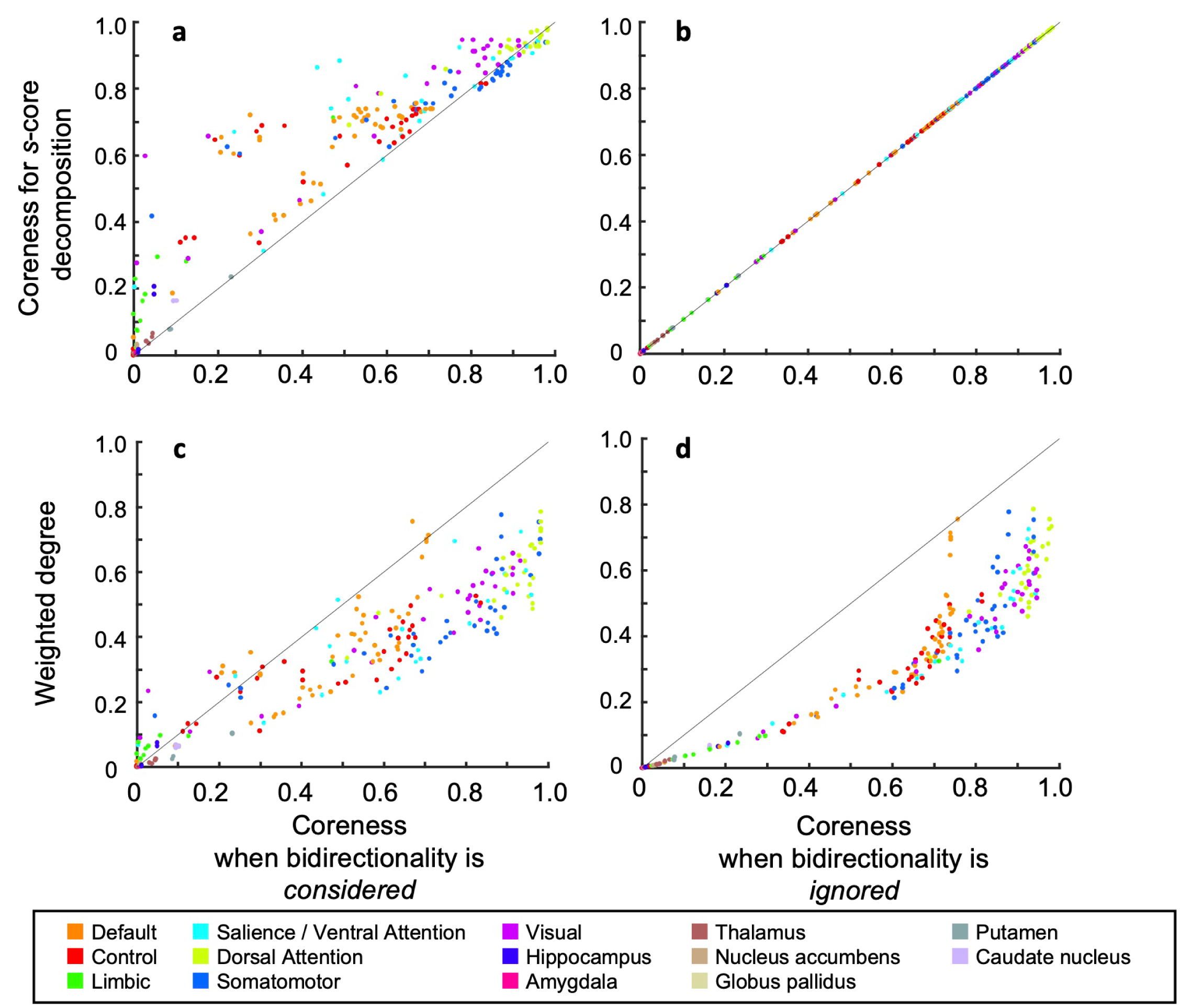
Comparison with other methods to extract strongly connected cores. Each metric is the average value across the resting state and seven tasks, after being normalized at the maximum in each brain state. Each point is color-coded according to the Yeo-7 network atlas and subcortical divisions. The solid line represents the identity line (*y* = *x*).

This means that the coreness for *s*-core decomposition demonstrated a weaker correlation with the MRR for iES or the functional connectivity gradient than did the coreness for the complexes when bidirectionality was considered.

Next, we compared with network hubs, which are nodes with a high degree. Here, the weighted degree of a node is defined as the sum of the weights of edges that are connected to the node, regardless of their directions (Eq. (14)). If strong complexes when bidirectionality is considered basically comprise hubs, this means that bidirectionality does not matter for the extraction of complexes. The results show that the coreness when bidirectionality was considered did not necessarily comprise hubs. Figure **7c** shows that strong complexes include not only hubs but also medium-degree nodes, although the weighted degree and the coreness exhibit a certain level of correspondence. Similarly, when bidirectionality is ignored, the coreness also shows correspondence with the weighted degree. However, there is a difference in that, particularly in regions with lower degrees, the weighted degree and coreness tend to match almost one-to-one (Fig. **7d**). Additionally, the weighted degree demonstrated a weaker correlation with the MRR and the functional connectivity gradient than did the coreness when bidirectionality was considered (MRR; *r* = 0.387 95% confidence interval (CI) [-0.087, 0.678], the central 95% interval of the null distribution based on brainSMASH [-0.577, 0.577], *p*_brainSMASH_ = 0.151, computed on *n* = 17 points, functional connectivity gradient; *r* = 0.277, 95% CI [0.149, 0.379], 95% null distribution interval (brainSMASH) [-0.315, 0.321], *p*_brainSMASH_ = 0.0529, *n* = 200 points, Extended Data Figs. **7-1a** and **7-1b**).

These results demonstrate that considering bidirectionality for extracting cores can reveal core structures of the human functional network that are unidentifiable when analyzed using simple methods that do not consider bidirectionality.

## Discussion

In this study, we investigated the subnetworks of the brain that exhibit strong or weak bidirectionality, and examined their relationship with perceptual awareness and other cognitive functions. To achieve this, we proposed a novel framework for extracting cores (complexes) with strong bidirectional interactions from brain activity (Fig. **1**). We applied this framework to HCP fMRI data of the resting state and seven cognitive tasks and extracted complexes. The analysis revealed the average tendencies of the regions included in the cores with strong and weak bidirectionality across different brain states, and how these are characterized in relation to perceptual awareness and other cognitive functions (Figs. **2**–**5**). Regarding the relationship with perceptual awareness, cerebral cortical regions were more likely to be included in the strongly bidirectional cores whereas subcortical regions were less likely to be included (Fig. **2**). Specifically within the cerebral cortex, there was a moderate positive correlation (*r* = 0.462) between iES-induced perceptual elicitation rates and the likelihood of being included in the strongly bidirectional cores (Fig. **3**), but the relationship is not statistically robust according to the 95% confidence interval and the p-value (95% confidence interval [-0.074, 0.755], central 95% interval of the null distribution [-0.583, 0.574], *p*_brainSMASH_ = 0.0863). It is important to note that the uncertainty is inherently large due to the small sample size of only *n* = 17 points. To pass the conventional threshold of *p*-value *<* 0.05, the correlation would have to be greater than *r* = 0.574, which is a considerably large correlation and generally difficult to achieve empirically. Additionally, the p-value obtained with brainSMASH is conservative because it preserves spatial autocorrelation. Taken together, although we cannot definitively conclude that there is a clear association due to the limited small sample size, we interpret the result of the positive correlation *r* = 0.462 as a non-negligible and moderate effect.

Regarding the relationship with other cognitive functions, a meta-analysis and a comparison with the functional connectivity gradient revealed that cores with stronger bidirectionality tended to be associated with lower-order, rather than higher-order, cognitive functions (Figs. **4** and **5**) (the correlation *r* = 0.397). It is worth noting that although the results with the functional gradient are statistically more robust (95% CI [0.299, 0.478], *p*_brainSMASH_ = 0.00280) than those with the mean response rate for iES, they are obtained based on a much larger sample size (*n* = 200) and thus, it is generally much easier to obtain a smaller p-value simply due to the larger sample size.

In the following subsections, we first explain the network structure revealed in this study using our framework for extracting subnetworks with strong bidirectional interactions. We then examine the relationship of cores that exhibit strong bidirectional interactions with conscious perception. Based on the results of this study, we argue that the regions with strong bidirectional interactions broadly correspond to regions considered important for conscious perception, although this relationship involves some uncertainty due to the small sample size. Subsequently, we discuss the underlying mechanisms of the observed potential correspondence between iES-induced perceptual elicitation rates and the strength of bidirectional interactions. Next, we discuss the interpretation of our findings suggesting that regions associated with lower-order sensorimotor processing belong more to strongly bidirectional cores compared to regions in the association cortices, which are known for their role in higher cognitive functions. Additionally, we contextualize our results by comparing the extracted cores with existing core extraction methods from a network analysis perspective. Finally, we discuss future prospects for applications of our proposed framework.

### The network structure revealed in this study

Our findings revealed that while cortical regions exhibit high coreness, subcortical regions show relatively lower coreness. This result suggests that, although strong bidirectional cores are formed within cortical regions, sub-cortical regions are not directly incorporated into these cores. A primary factor contributing to this phenomenon is the relatively weaker statistical causal strength within subcortical regions and between subcortical and cortical regions, as illustrated in the constructed directed graph (Extended Data Fig. **2-1a**).

### Correspondence between the strength of bidirectional interactions and the significance for conscious perception

In this subsection, we discuss a potential link between a region’s inclusion in the central cores and its importance for conscious perception. This argument is supported by two key findings: firstly, cortical regions are identified as more likely to be included in central cores than subcortical ones (Fig. **2**). Secondly, there is a moderate positive correlation between the coreness and the iES-induced perceptual elicitation rates, although this finding is not statistically robust due to the small sample size. (Fig. **3**).

Regarding the first finding, while the tendency for cortical regions to show higher coreness likely reflects the importance of the cerebral cortex in conscious perception, the interpretation of the tendency for subcortical regions to show lower coreness remains less clear. The significance of the cerebral cortex in conscious perception has been well-established through previous research (Koch et al., 2016; Lamme, 2018; Mashour et al., 2020; Marshel et al., 2019; Filipchuk et al., 2022). Therefore, it can be said that the observation that cortical regions tend to be included in the central cores (i.e., exhibit higher coreness) aligns with this significance. On the other hand, while some studies argue that subcortical regions play a less direct role in conscious perception (Koch et al., 2016; Lamme, 2018; Mashour et al., 2020), others claim they are necessary (e.g., (Aru et al., 2019; Ward, 2011; Slagter et al., 2017; Afrasiabi et al., 2021), brainstem; (Edlow et al., 2024), thalamus; (Whyte et al., 2024)). Therefore, further careful research is necessary to determine whether the lower coreness observed in subcortical regions corresponds to the actual contribution of subcortical regions to conscious experience.

Regarding the second finding, the relationship between iES-induced perceptual elicitation rates and regions with high coreness suggests that these core areas may play an important role in conscious perception. iES allows for the causal modulation of neural activity, indicating that stimulation of a specific region leading to changes in conscious perception demonstrates the direct involvement of that region in that perception (Raccah et al., 2021). Therefore, the observed correlation between coreness and perceptual elicitation rates implies that central core regions within the cerebral cortex may be important for conscious perception.

Nevertheless, this comparison must be interpreted with caution due to several limitations. iES can also induce motor responses, raising the possibility that changes in conscious perception may result from awareness of the induced movement rather than the direct involvement of the stimulated region. Additionally, it is important to recognize that this comparison is restricted to brain regions responsible for perceptual awareness within the range of perceptual categories elicited by iES experiments (see Methods for details).

Overall, while the findings suggest a potential link between regions included in the bidirectional cores and their importance for conscious perception, we cannot draw definitive conclusions due to the various factors mentioned above. Further research is needed to clarify the relationship between strongly bidirectional cores and conscious perception.

### Interpretation of the mechanism behind the high elicitation rates induced by iES for regions within cores

Below, we discuss the underlying mechanism behind a potential positive correlation between high elicitation rates induced by iES for regions and strongly bidirectional cores. As regions within a core are strongly connected to other regions within the core, it is expected that stimulating a region within the core will propagate its effects throughout the core, and thereby result in a change in conscious perception. Our results do not contradict this expected mechanism, supporting the observed general tendency for the high elicitation rates induced by iES for regions within cores. On the other hand, it is also expected that even when a region outside a core is stimulated, if the region provides input to the core, the effect of the stimulus could propagate to the core, and thus result in a change in perception. However, our results showed that the perceptual elicitation rates in regions not included in the strongly bidirectional core were generally low (Fig. **3**, Extended Data Figs. **6-1**). This indicates that when a region that causally affects the core, that is, provides input to the core, is stimulated, there tends to be a lower tendency to induce changes in perception. Whether a perceptual change occurs when a region outside the core is stimulated depends on factors such as the intensity of the stimulation. Even when stimulating non-core regions that provide input to the core, perceptual change could occur if the stimulus intensity is strong enough.

To validate this interpretation in the future, it will be crucial to monitor the extent and manner in which the signal propagates bidirectionally upon stimulation. Additionally, it will be important to determine whether the resulting perception aligns with the perception associated with the region to which the signal has propagated. This will involve adjusting the intensity of the stimulation and correlating it with the outcomes of extracted cores. Such analysis requires measurement techniques with high temporal resolution that are capable of capturing signal propagation and perceptual changes within a timeframe of milliseconds to a few seconds. However, the temporal resolution of fMRI exceeds this timeframe, being longer than a few seconds (Glover, 2011). Therefore, incorporating EEG or ECoG data in future studies could be beneficial, offering higher temporal resolution than fMRI. These methods enable the capturing of rapid signal dynamics and perception changes, effectively complementing fMRI’s spatial insights.

### Correspondence of the strength of bidirectional interactions with other cognitive functions

In this study, a meta-analysis using NeuroSynth and a comparison with the functional connectivity gradient revealed that cores with strong bidirectional interactions tended to include regions associated with lower-order sensorimotor functions rather than regions associated with higher-order cognitive functions. This section elucidates the implications of these findings and explores potential explanations for these observations.

The results in this paper suggest that lower-order regions form cores in which they interact with each other in a bidirectional manner, whereas other regions have weak bidirectional interactions with those cores. Lower-order sensory processing regions are the first in the cortex to receive stimuli from the outside world and play a fundamental role in sending signals to higher-order regions. Lower-order regions also receive top-down signals from higher-order regions. This suggests that lower-order regions function as the cores of multidirectional information flow by coordinating with each other.

Next, we discuss the factors that led to the result that lower-order regions form cores with strong bidirectional interactions. The simplest, albeit naïve, interpretation is that this result is due to differences in the strength of bidirectional interactions. That is, the interpretation is that the bidirectional interactions are stronger between lower-order regions, while such interactions are weaker between lower-order and higher-order regions and between higher-order regions by comparison.

Another interpretation is that the present results were obtained because bidirectional interactions universally occur in lower-order regions, but whether such interactions occur between lower- and higher-order regions depends on many factors, including attention (Gregoriou et al., 2009; Baldauf and Desimone, 2014) and strength of sensory stimuli (van Vugt et al., 2018). It should also be noted that this study analyzed average trends across resting and seven different task states (Van Essen et al., 2013; Barch et al., 2013). It is possible that in lower-order regions, task-independent bidirectional interactions occur, while in higher-order regions, they may be task-dependent.

To better understand the relationship between cognitive functions and bidirectional interactions, it will be important to analyze data that takes into account the abovementioned factors, as well as a more detailed analysis of task-specific brain states.

### Comparison with other core extraction methods in terms of association with cognitive functions

In this research, we evaluated our proposed methodology against traditional core extraction techniques, namely *s*-core decomposition and network hubs, which do not consider bidirectionality (Fig. **7**). In what follows, we compare these methods in the context of their association with cognitive functions.

Our findings revealed an association between the coreness of our method and two key metrics: the MRR of iES and the functional connectivity gradient. Conversely, the *s*-cores and hubs demonstrated a weaker correlation with these metrics. This suggests that our approach can unveil connections between network cores and cognitive functions that remain undetected by conventional methods due to their non-consideration of bidirectionality.

Unlike our study, previous research utilizing *s*-core decomposition and network hubs have identified central cores or hubs primarily identified network cores within higher-order regions (Achard et al., 2006; Achard and Bullmore, 2007; Buckner et al., 2009). In contrast, our study revealed that the central cores and hubs were primarily situated in lower-order regions. This divergence likely stems from our approach of employing normalized directed transfer entropy (NDTE, (Deco et al., 2021)) as an edge weighting metric. NDTE captures the directional and statistical causal relationships within the network, unlike previous studies that relied on correlations as edge weights, ignoring the directionality or statistical causality of interactions. This methodological difference elucidates the distinct outcomes between our findings and those of previous research, highlighting the significance of considering directionality in network analysis.

In summary, our research underscores the value of incorporating directionality based on estimated directed influences in network analysis. Through this innovative approach, we have established a new link between network cores with strong bidirectional interactions and crucial neurological metrics, namely the MRR and the functional connectivity gradient, enhancing our understanding of the relationship between the brain network structure and its functions.

### Distinguishing features of our framework compared to research proposing the adopted graph construction method

From the perspective of core extraction for directed networks, our framework for extracting cores in brain-directed networks has unique characteristics in both the methodology for core extraction and the regions identified as cores. However, it is worth mentioning that our framework adopts the graph construction method used by Deco et al. (Deco et al., 2021), indicating that our framework does not introduce novelty in the graph construction process. Instead, the extraction of bidirectional cores serves as a key element that characterizes our framework. Notably, while they also extract core regions from directed networks, their approach differs from ours. Of course, in addition to the fundamental differences in research objectives between our study and theirs, the analyses, such as the comparison of coreness with iES or functional gradients, are also decisively different. Along with these fundamental differences, the methodologies and regions extracted as cores differ between the two studies. Given that both studies share the same graph construction method but adopt different core extraction techniques, the differences in core extraction methodology and extracted regions can be regarded as elements that characterize our framework from the perspective of core extraction. Therefore, in the following section, we explain the distinguishing aspects of our core extraction method in comparison with the study by Deco et al. First, we will explain the methodological features of our core extraction method, followed by a discussion of the regions identified as cores.

First, compared to a wide range of existing core extraction methods, our method has unique characteristics—globality, bidirectionality, and exactness—which also apply to the study by Deco et al. (See Methods for details). The first distinct feature of our method is that it extracts cores that are densely bidirectionally connected. Considering that bidirectional interactions are regarded as crucial for conscious perception, this capability is especially important in this context of neuroscience. While Deco et al.’s approach extracts densely connected subnetworks, it does not impose the requirement for bidirectional connections within those cores. As an illustrative example, consider the graph in Extended Data Fig. **1-1a** and Extended Data Figure. **1-2a**. In this case, our method extracts only the bidirectionally connected subnetwork as the most central core (Extended Data Fig. **1-1a**, a node set {EFIJ}). In contrast, the functional rich club method extracts a network that includes nodes that are not bidirectionally connected (e.g., nodes A, D, H) as the most central core (Extended Data Figure. **1-2a**, a node set {ABDEFHIJ}).

The second distinction is that our method extracts cores by considering the global structure of the network, taking into account whether nodes are interconnected across the entire network, whereas the functional rich club method extracts cores based on local metrics and cannot account for such network-wide connectivity. One example illustrating this difference is the case where two modules are connected by bidirectional edges (Extended Data Fig. **1-1c****, d**). In this scenario, when considering the global structure, two cores would be expected to be extracted. Indeed, in such cases, our method identifies two sub-networks as the most central cores (see Extended Data Fig. **1-1c**), whereas the functional rich club method is likely to identify the entire network as the most central core (see Extended Data Fig. **1-1d**).

The final distinct feature of our method is the exactness of core extraction, whereas other methods are based on approximation. Due to computational constraints, Deco et al. apply approximation for core extraction in cases with a large number of nodes. By contrast, our method leverages a fast core extraction algorithm (Kitazono et al., 2023), allowing us to perform exact core extraction even in cases with a large number of nodes.

The methodological differences in core extraction lead to differences between the functional rich club method and our core extraction method in terms of the identified core regions. Specifically, the functional rich club method identifies cortical regions such as the precuneus and the posterior and isthmus cingulate cortex, which are part of the Default Mode Network, as central cores, as well as subcortical regions like the hippocampus. On the other hand, our core extraction method identifies unimodal sensory processing regions as central cores.

### Future directions

The present study targeted human subjects; nevertheless, the extension of future analyses to include non-human species is an intriguing possibility (Xu et al., 2020; Goulas et al., 2014; Eichert et al., 2020; Fulcher et al., 2019). Experimental evidence indicates that bidirectional interactions play a pivotal role in conscious perception across various species (Lamme et al., 1998; Supèr et al., 2001; Cauller and Kulics, 1988; Cauller and Kulics, 1991; Self et al., 2012; Koivisto et al., 2014; Sachidhanandam et al., 2013; Manita et al., 2015; Nieder et al., 2020; Cohen et al., 2018). A comparative study investigating whether the cores with strong bidirectional interactions in non-human species consist of regions analogous to those that were identified in the strong cores in this study could significantly enhance our understanding of the relationship between bidirectional interactions and conscious perception. Such an exploration could offer a broader perspective on the neural mechanisms underlying conscious perception across different species and shed light on the evolutionary aspects of consciousness. Cross-species comparison is also interesting for understanding the relationship between the cores and lower- and higher-order cognitive functions. Lower-order cognitive functions, such as sensory perception, are common to many species, whereas higher-order cognitive functions are more developed in more advanced species. By analyzing cores with bidirectional interactions and their correlation with both lower- and higher-order cognitive functions in different species, we can elucidate the fundamental impact of bidirectional interactions on cognitive functionality.

The current study delved into the association between cores with strong bidirectional interactions and conscious perception; however, further investigation into how cores correlate with different consciousness states— such as wakefulness, sleep, anesthesia, and coma—is required. Previous research has highlighted the role of bidirectional interactions not only in the generation of conscious perception but also in maintaining wakefulness (Tononi et al., 2016). Additionally, recent studies revealed that the degree of system-level integration between brain regions was associated with consciousness states (Luppi et al., 2019; Luppi et al., 2021; Onoda and Akama, 2023). In light of these considerations, a crucial next step would involve identifying cores from datasets acquired during unconscious states, such as sleep or anesthesia, and contrasting these with observations from wakeful states. Such a comparison could provide deeper insights into the relationship between bidirectional interactions and varying levels of consciousness and enhance our understanding of the neurobiological mechanisms governing consciousness and its different states.

## Acknowledgement

This study was supported by Graduate Program for Social ICT Global Creative Leaders of the University of Tokyo by MEXT to TT, JST ACT-X Grant Number JPMJAX20A6 to JK, JST Moonshot R&D Grant Number JPMJMS2012 to SS and MO, JST CREST Grant Number JPMJCR1864 to MO, and JSPS KAKENHI Grant Numbers 18H02713 and 20H05712 to MO. Data were provided in part by the Human Connectome Project, WU-Minn Consortium (Principal Investigators: David Van Essen and Kamil Ugurbil; 1U54MH091657) funded by the 16 NIH Institutes and Centers that support the NIH Blueprint for Neuroscience Research; and by the McDonnell Center for Systems Neuroscience at Washington University. We acknowledge the use of ChatGPT (Open AI, https://chat.openai.com) to proofread our draft. We would like to thank Editage (www.editage.jp) for English language editing.

## Extended Data Legends

**Figure 1-1:**
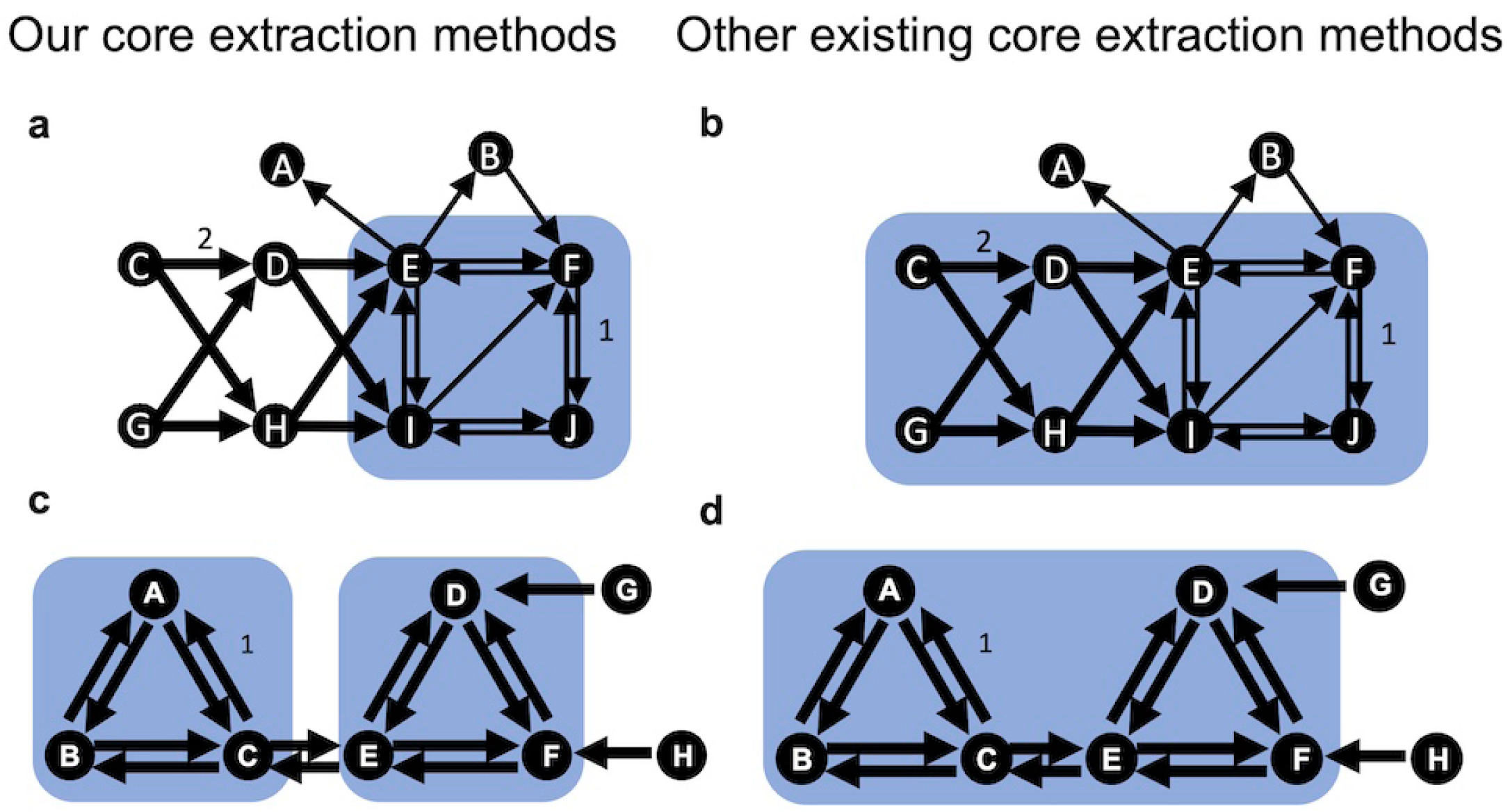
Examples of the differences between our core extraction method and other core extraction methods. **a, b**, Example of a graph *W* illustrating the difference in bidirectionality of the extracted cores. This is the same as Figure. **1d**. In this case, our method extracts only the bidirectionally connected subnetwork (a node set {EFIJ}) as the most central core (**a**), whereas s-core decomposition extracts a subnetwork that include unidirectionally connected nodes (e.g., nodes C, D, G, H) as the most central core (**b**, a node set {CDEFGHIJ}). In the case of s-core decomposition, it was applied to the symmetrized undirected network (*W* + *W^T^*)*/*2. **c, d**, Example illustrating the difference in globality of core extraction. In this network, two modules (a sub-graph containing nodes A, B, C and that containing nodes C, D, E) are connected by bidirectional edges. The weight of all edges is set to 1. In this case, other existing core extraction methods (such as s-core/k-core decomposition) extract a single, nearly entire network as the most central core (**d**), whereas our method extracts two sub-networks as the most central cores (**c**).

**Figure 1-2:**
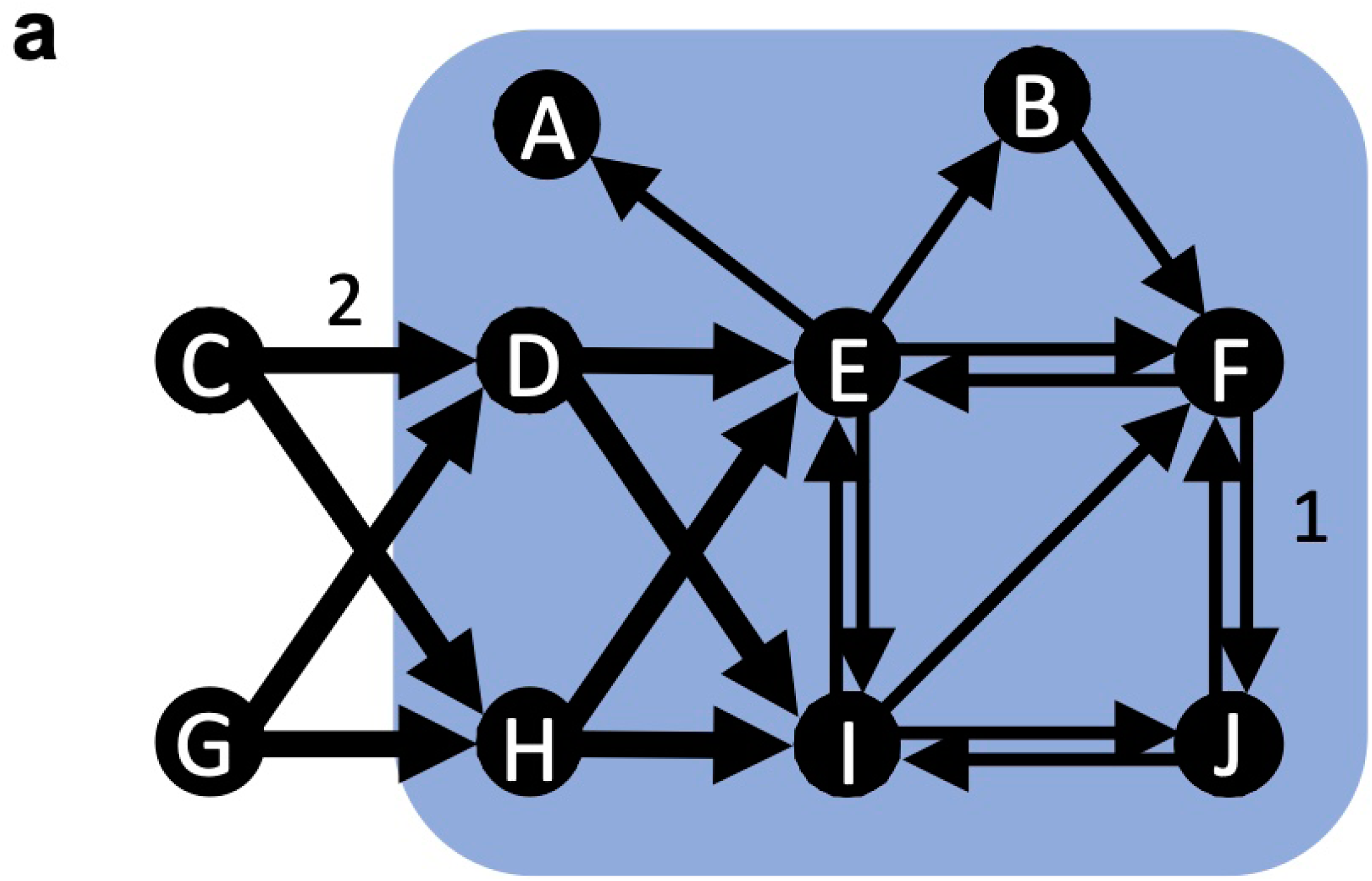
An example of extracted cores by the functional rich club (Deco et al., 2021). **a**, The network is the same as Figure. **1d**. The functional rich club method extracts a subnetwork that include unidirectionally connected nodes (e.g., node A,D,H) as the most central core (**b**, a node set {ABDEFHIJ}). Here, *G*_FRIC_(*k*) is defined as *G*_FRIC_(*k*) = Σ*_i_*_∈_*_k_* Σ*_j_*_∈_*_k_ W_ij_* + Σ*_i_*_∈_*_k_* Σ*_j_ W_ij_* − Σ*_i_* Σ*_j_*_∈_*_k_ W_ij_* (where *k* is a subset of regions *k* = {*i*_1_, *i*_2_, *…, i_l_*}), and the core was identified as the largest subnetwork among those whose *G*_FRIC_(*k*) fall within the top 5% of subnetworks of the same size.

**Figure 2-1:**
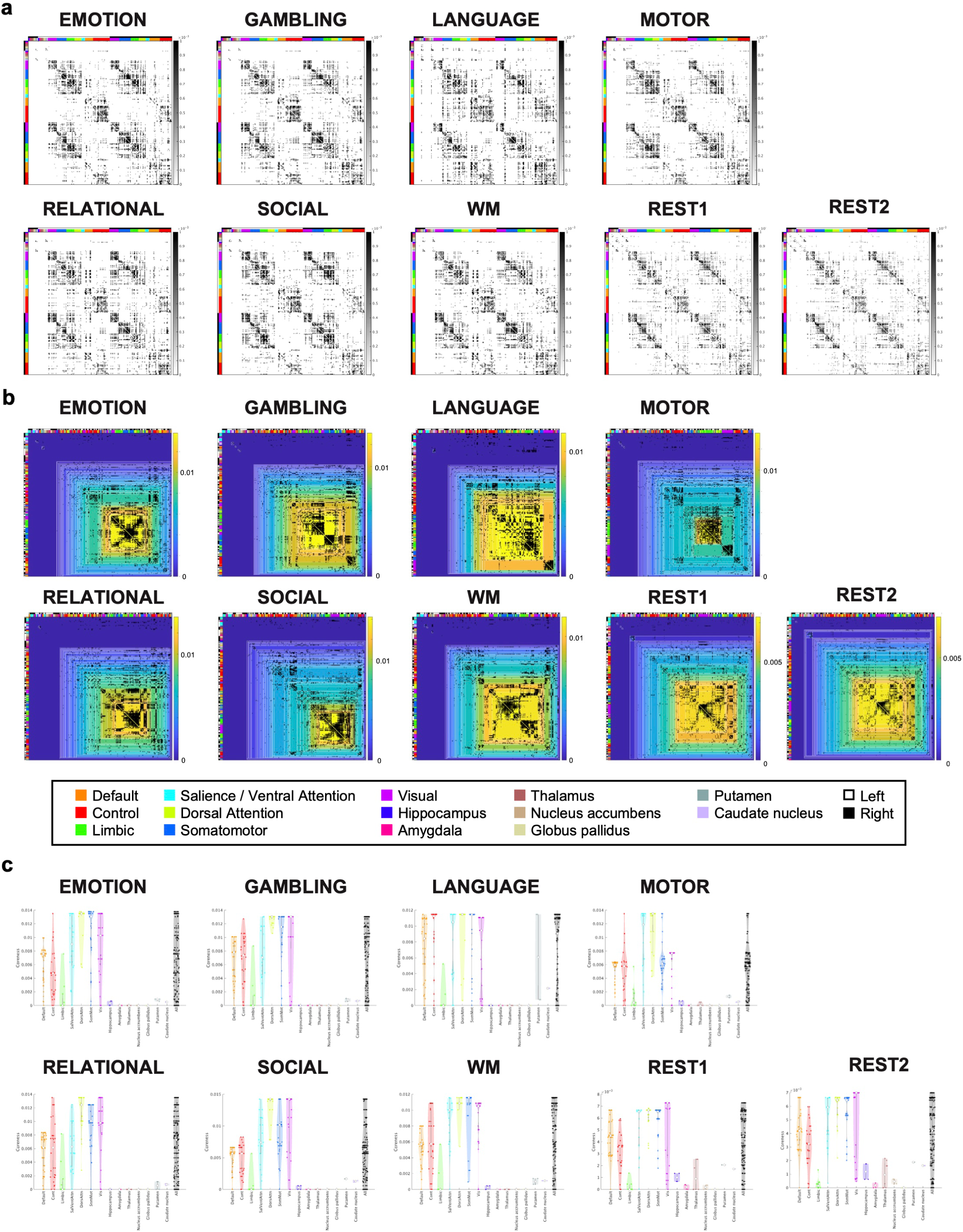
Results of the bidirectional cores extracted for each task. **a**, Estimated directed graphs at rest and during seven tasks. The grayscale of each matrix element represents the weight of the edge in the estimated directed network. The white and black at the left and top of the matrix indicate the left and right hemispheres, respectively, whereas the color bar represents the divisions of the Yeo-7 network atlas and major divisions of subcortical areas. **b**, Hierarchical structure of the cores with bidirectional connections at rest and during seven tasks. The directed network is arranged according to the hierarchical structure of the complexes, with white lines separating each hierarchical level. **c**, Violin plots showing the distribution of coreness at rest and during seven different tasks, categorized by the Yeo-7 network atlas for cortical regions and major divisions for subcortical areas. **a, b, c** show, from left to right, the results for emotion, gambling, language, motor, relational, social, working memory, rest1 and rest2, respectively.

**Figure 2-2:**
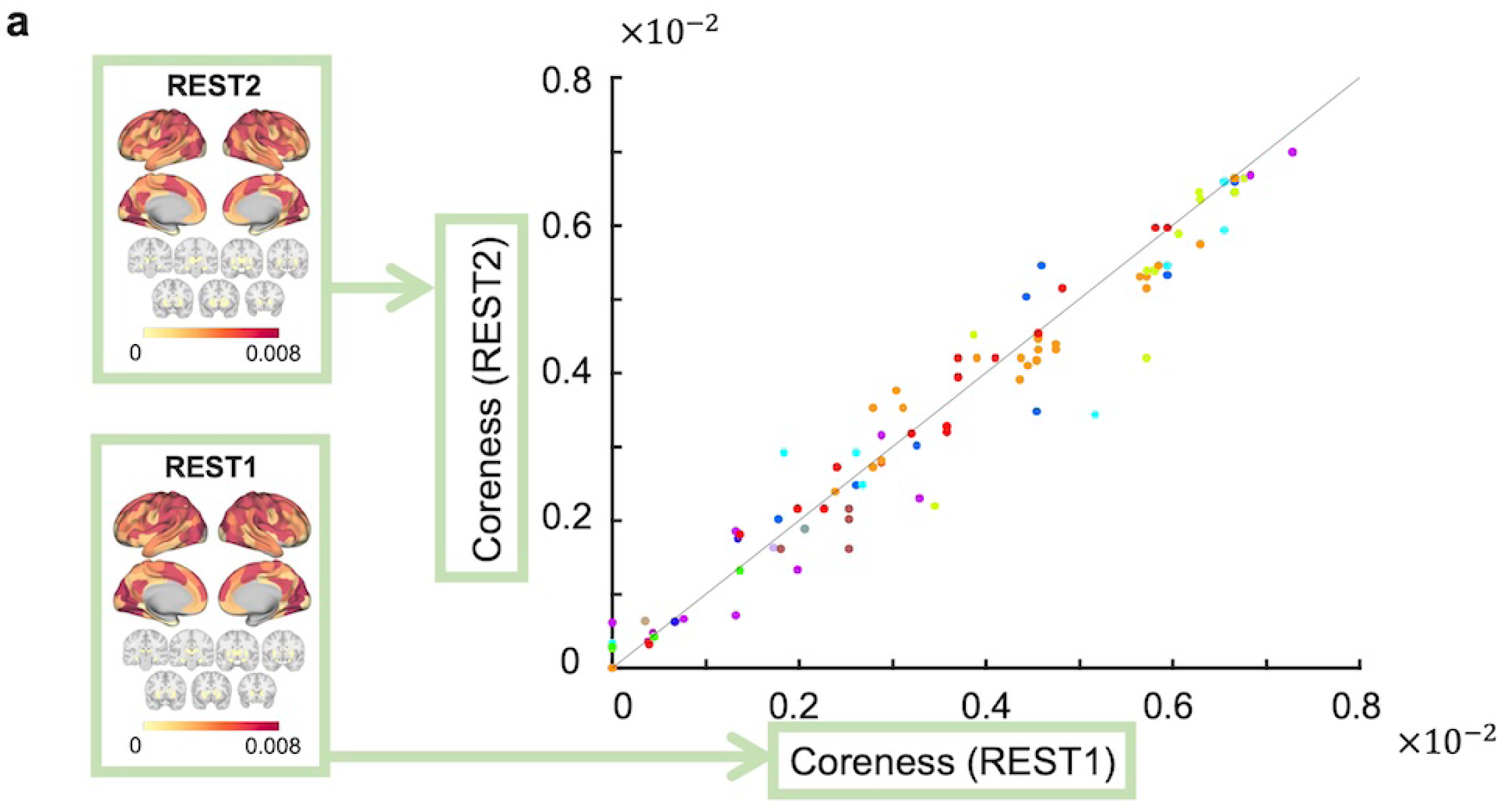
The similarity of coreness between Rest 1 and Rest 2. **a**, (Left:) Coreness at Rest 1 (top) and Rest 2 (bottom) in the cerebral cortex and the subcortex shown in seven coronal slices. (Right:) A scatterplot of coreness at Rest 1 and Rest 2. The solid line represents the identity line (*y* = *x*).

**Figure 2-3:**
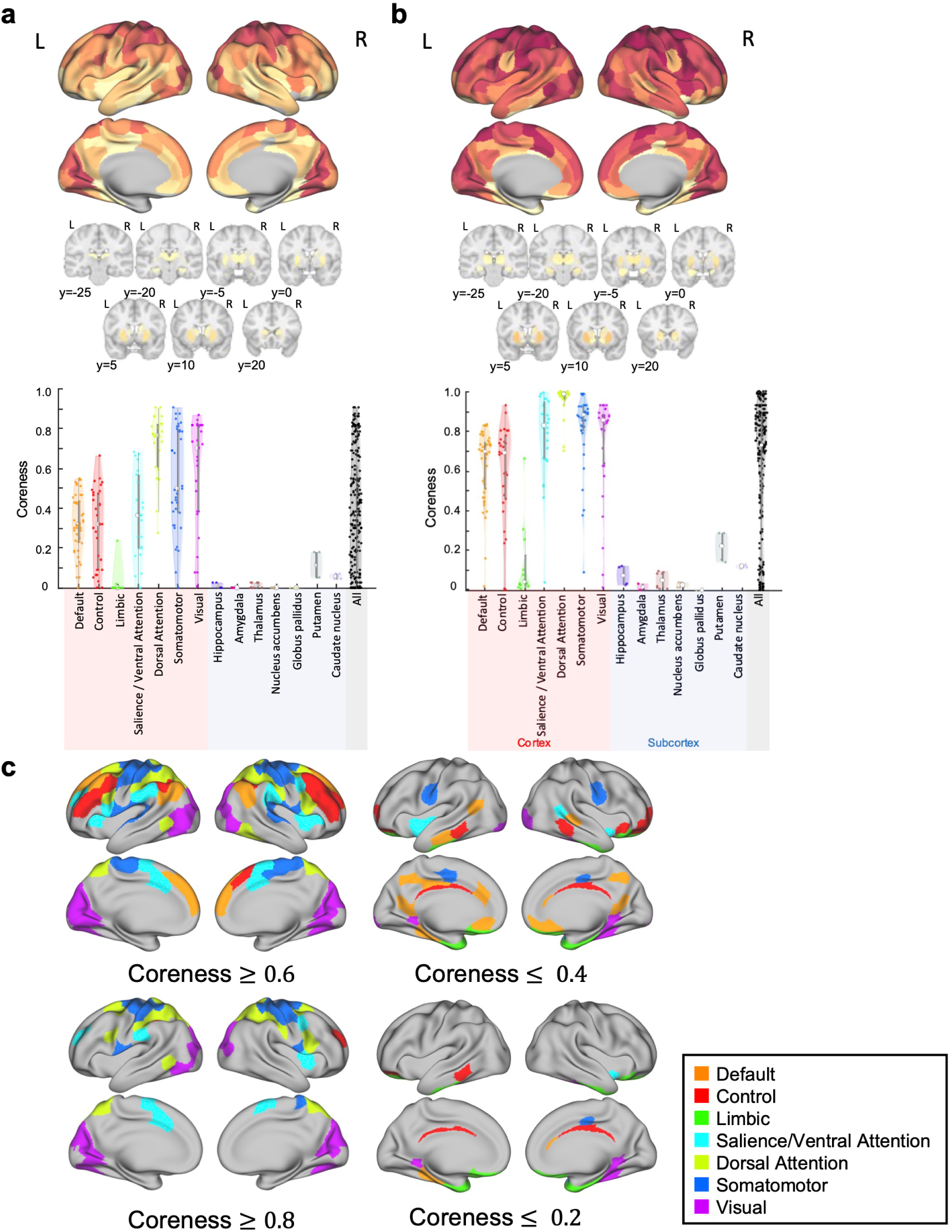
The robustness of trends in the extracted bidirectional cores when graph density or thresholding value of coreness is varied. **a, b**, Coreness at 5% (**a**) and 20% (**b**) graph density in the cerebral cortex (top) and the subcortex shown in seven coronal slices (middle), along with violin plots of coreness (bottom). **c**, ROIs with coreness greater than 0.6 or 0.8, or less than 0.4 or 0.3, are colored according to the Yeo-7 network atlas.

**Figure 3-1:**
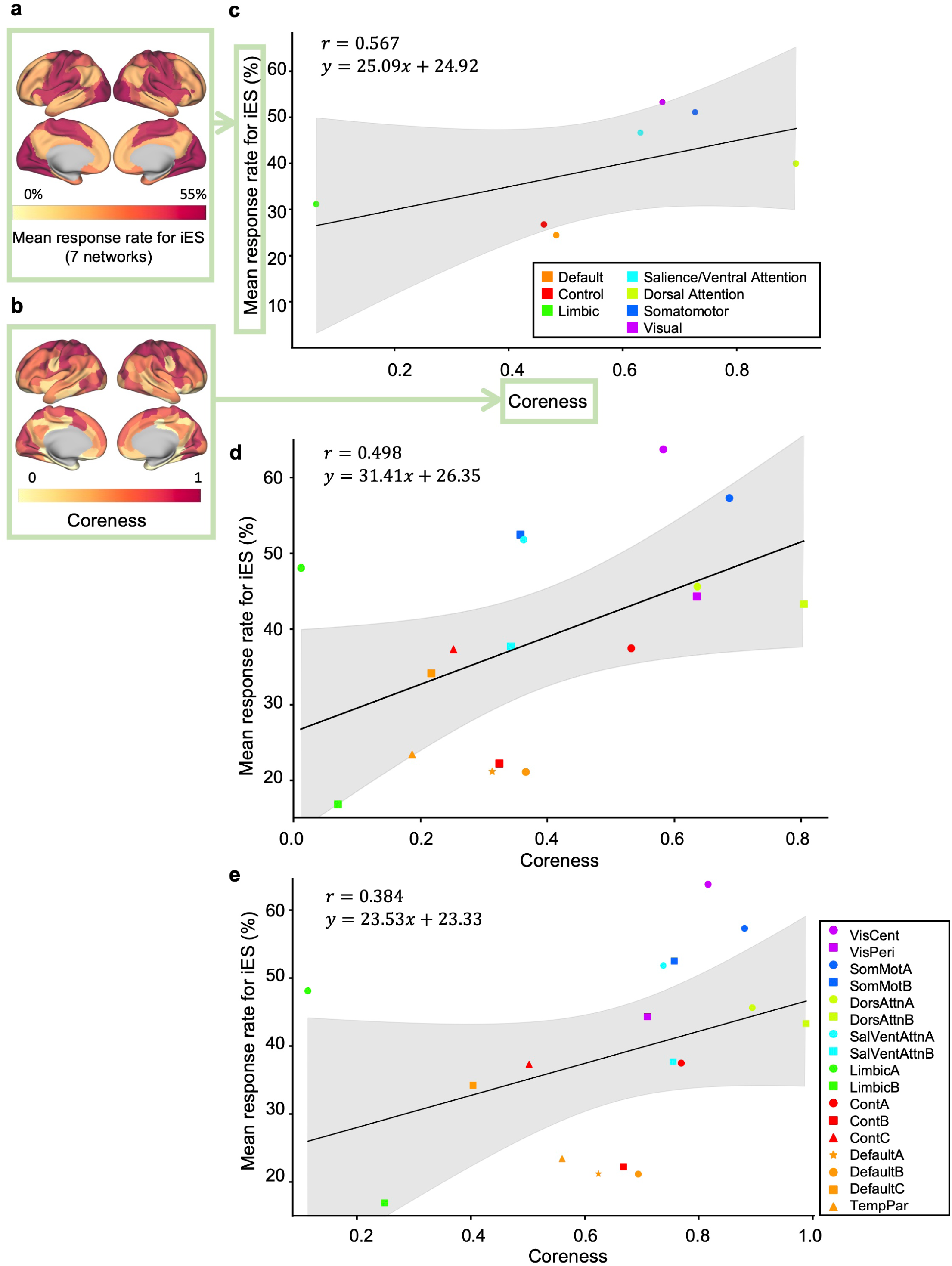
The robustness of the trend of positive correlation between coreness and MRR when Yeo-7 network atlas is used and graph density is varied. **a, b**, A cortical surface rendering of the MRR for iES (7 network) (**a**), and that of coreness (**b**). **c**, A scatter plot showing the average coreness for each division of the Yeo-7 network atlas (horizontal axis) and the MRR (vertical axis). The solid line represents the regression line. Each point is color-coded according to the Yeo-7 network atlas. **d, e**, Scatter plots showing the average coreness for each division of the Yeo-17 network atlas (horizontal axis) and the MRR (vertical axis) at 5% graph density (**d**) and 20% graph density (**e**).

**Figure 4-1:**
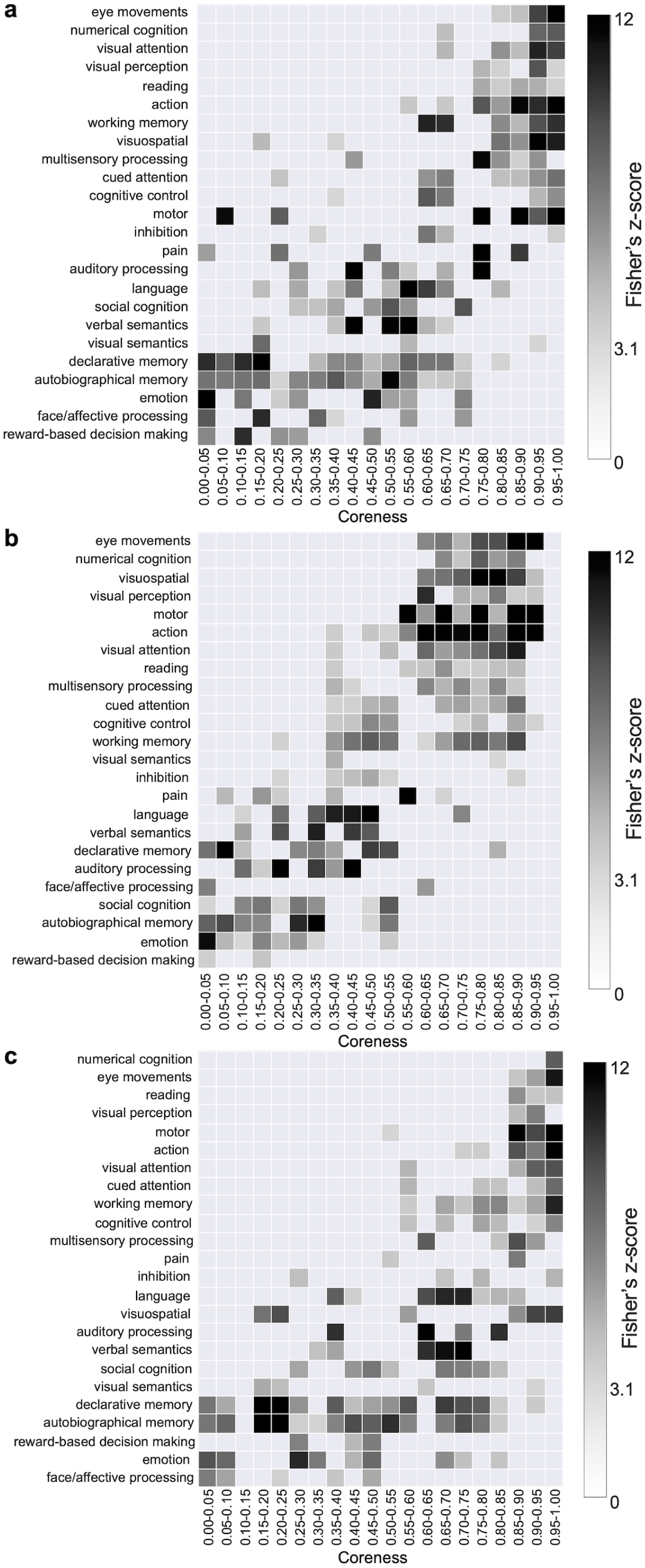
The robustness of the observed relationship found in the Term-based meta-analysis of the core structure using NeuroSynth when subcortical regions were included and graph density was varied. **a, b, c**, A term-based meta-analysis using NeuroSynth applied to coreness when subcortical regions were included (**a**), and when graph density was 5% (**b**) and 20% (**c**).

**Figure 5-1:**
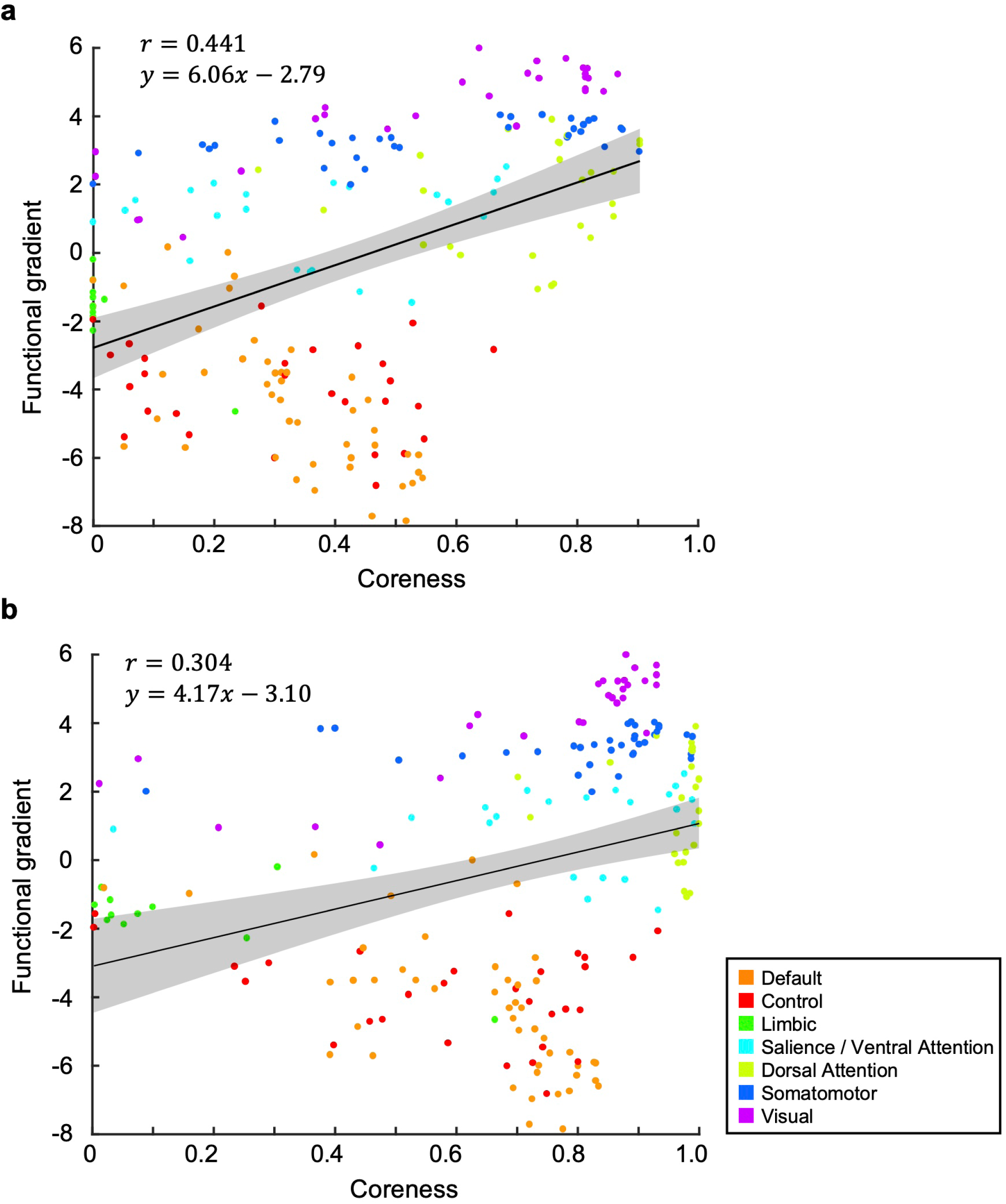
The stability of the positive correlation between coreness and the functional connectivity gradient when graph density was varied. **a, b**, Scatterplots of coreness and the functional connectivity gradient when graph density was 5% (**a**) and 20% (**b**).

**Figure 6-1:**
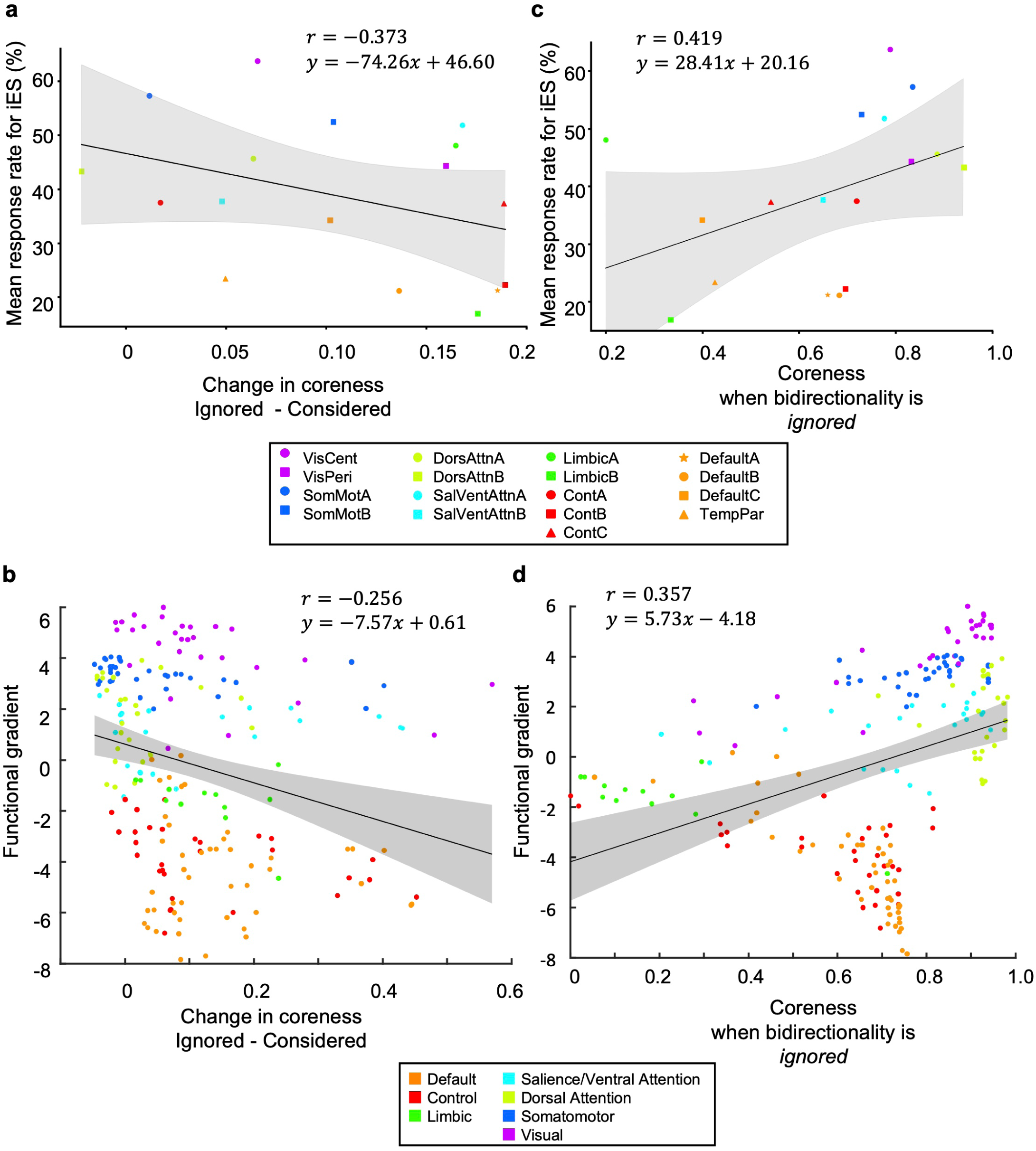
Comparison of changes in coreness when ignoring bidirectionality and coreness when ignoring bidirectionality with the MRR for iES and the functional connectivity gradient. **a**, There is a negative correlation between the change in coreness and the MRR for iES (*r* = −0.373). **b**, There is a negative correlation between the change in coreness and the functional connectivity gradient (*r* = −0.256). The solid lines represent the regression lines. **c**, Compared to when bidirectionality is considered, the correlation between coreness and the MRR is weaker when bidirectionality is ignored (*r* = 0.419). **d**, Compared to when bidirectionality is considered, the correlation between coreness and the functional connectivity gradient is weaker when bidirectionality is ignored (*r* = 0.357). The solid lines represent the regression lines.

**Figure 7-1:**
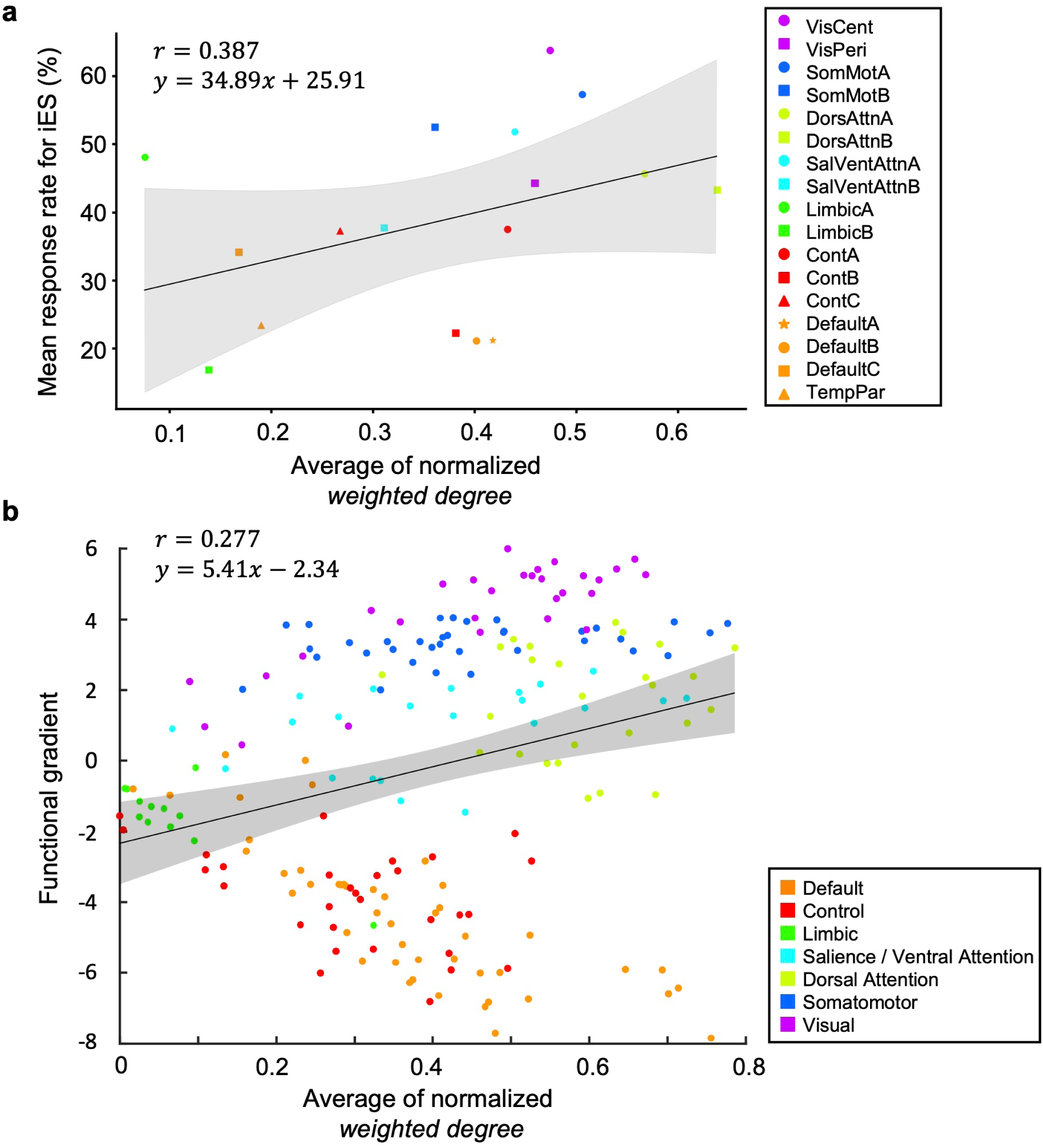
Comparison of weighted degree with the MRR for iES and the functional connectivity gradient. **a**, The correlation between the weighted degree and the MRR (*r* = 0.387) is weaker than that between the coreness when bidirectionality is considered and the MRR. **b**, The correlation between the weighted degree and the functional connectivity gradient (*r* = 0.277) is weaker than that between coreness when considering bidirectionality and the functional connectivity gradient. The solid lines represent the regression lines.

## Notes

**Financial Interests of Conflicts of Interests** The authors declare no competing interests.

### Competing Interest Statement

The authors have declared no competing interest.

### Summary of Updates

Discussion added; Figures revised; Statistical analysis in the Methods section revised.

